# ADAR1 haploinsufficiency and sustained picornaviral RdRp dsRNA synthesis synergize to dysregulate RNA editing and cause multi-system interferonopathy

**DOI:** 10.1101/2025.01.21.634124

**Authors:** Caitlin M. Miller, James H. Morrison, Laura Bankers, Rachael Dran, Julia M. Kendrick, Emma Briggs, Virginia L. Ferguson, Eric M. Poeschla

**Affiliations:** Division of Infectious Diseases, Anschutz Medical Campus, University of Colorado School of Medicine, Aurora, Colorado, USA 80045; Department of Mechanical Engineering and BioFrontiers Institute, University of Colorado at Boulder, Boulder, CO, 80309

**Keywords:** ADAR1, MDA5, innate antiviral immunity, picornavirus, RdRp, autoimmunity, autoinflammation, interferonopathy, lupus

## Abstract

Sensing of viral double-stranded RNA by MDA5 triggers abundant but transient interferon-stimulated gene (ISGs) expression. If dsRNA synthesis is made persistent by transgenically expressing a picornaviral RNA-dependent RNA polymerase (RdRp) in mice, lifelong MDA5-MAVS pathway activation and marked, global ISG upregulation result. This confers robust protection from viral diseases but in contrast to numerous other chronic MDA5 hyperactivation states, the mice suffer no autoimmune or other health consequences. Here we find they further confound expectations by being resistant to a strong autoimmunity (lupus) provocation. However, knockout of one allele of *Adar* breaks the autoinflammation-protected state of RdRp^tg^ mice and results in a severe disease that resembles interferonopathies caused by MDA5 gain-of-function protein mutations. *Adar*^+/-^ mice are healthy but *Adar*^+/-^ RdRp^tg^ mice have shortened lifespan, stunted growth, premature fur graying, poorly developed teeth, skeletal abnormalities, and extreme ISG elevations. A-to-I edits are both abnormally distributed and increased (numbers of genes and sites). These results, with a nucleic acid-triggered and MDA5-wild type model, illuminate the ADAR1-MDA5 axis in the regulation of innate immunity and establish that viral polymerase-sourced dsRNA can drive autoinflammatory disease pathogenesis.

**IMPORTANCE:** RNA virus double-stranded RNAs are important pathogen associated molecular patterns that are sensed by the RIG-I like receptor MDA5, which triggers an acute innate immune response involving many ISGs. One key to a healthy innate immune system is that MDA5 not sense endogenous dsRNA. This is normally ensured by dsRNA duplex-disrupting ADAR1 editing of host dsRNAs. Picornavirus RdRp^tg^ mice have an unusual constitutive MDA5 activation state, with very high lifelong MDA5-mediated ISG expression that confers robust protection from diverse lethal viruses. Importantly, and in contrast to numerous other chronic MDA5 hyperactivation states, the mice develop no autoinflammatory consequences. If we delete one ADAR1 allele, however, which by itself is well tolerated, the mice develop a multisystem disease that resembles the human interferonopathy Singleton-Merten syndrome. In contrast to other MDA5/ADAR1 disease models, the MDA5 and ADAR1 proteins are both wild type in this dsRNA-driven model.

## INTRODUCTION

Double-stranded RNA (dsRNA), a requisite intermediate during the replication of RNA viruses, is a pathogen-associated molecular pattern that induces an array of first-line antiviral defenses. Two specialized RNA helicases function as the main pattern recognition receptors (PRRs) that detect viral dsRNA: retinoic acid inducible gene-I (RIG-I) and melanoma differentiation antigen 5 (MDA5). Both of these RIG-I-like receptors (RLRs) are cytosolic sensors that signal through the adaptor MAVS, which instigates signaling cascades that eventuate in expression of numerous ISGs, either directly or via induction of secreted type I interferons (IFN-I) (1). RIG-I in general detects shorter or 5’-ppp containing dsRNAs and MDA5 detects the internal segments of longer (> 1 to 2 kb) dsRNAs (2–4). A few RNA viruses are exclusively sensed by one RLR (1). Picornaviral dsRNAs, for example, are detected by MDA5 and not RIG-I (1, 5–8).

The challenge such a system poses for the host is to securely differentiate viral dsRNAs from the vast, diverse pool of cellular RNAs, many of which can also harbor extended RNA duplex segments, particularly within retroelement-derived RNAs that are abundant in mammalian genomes (9, 10). Inappropriate or sustained activations of RLRs and other signaling pathways by endogenous nucleic acid ligands are associated with a variety of autoimmune diseases, which include classical, common syndromes such as systemic lupus erythematosus (SLE), type I diabetes mellitus, and psoriasis as well as various and frequently severe genetic conditions known collectively as interferonopathies. These auto-inflammatory conditions are characterized by persistent IFN-I and ISG expression and diverse end organ pathologies (5, 11, 12).

The RNA modifying enzyme Adenosine Deaminase Acting on RNA 1 (ADAR1) has recently been identified as a central constraint on dsRNA-triggered autoimmunity (13–17). Encoded by *ADAR*, ADAR1 catalyzes post-transcriptional editing of host RNAs by deaminating adenosine to inosine, thereby disrupting A-U base-paring and preventing formation of long uninterrupted RNA duplexes (9, 10, 18, 19). A-to-I editing most prominently prevents detection by MDA5 or protein kinase regulated by dsRNA (PKR) of endogenous retroelement transcripts and other dsRNAs, which averts inappropriate immune activation or translational shutoff (9, 10). Such editing of transcripts can also cause protein recoding (17, 20, 21), although the vast majority of A-to-I edits in mammals are in non-coding regions (22–25). Strikingly, physiologically essential A-to-I editing represents a very small fraction of the editome, and moreover most editing is unnecessary for murine homeostasis in the absence of MDA5 (26, 27). Hereditary mutations in *ADAR*, as well as in other genes such as *TREX1*, *SAMHD1*, *RNASEH2, RNU7, LSM11, and IFIH1* (MDA5) have been linked to the development of the rare congenital inflammatory disorder Aicardi*-*Goutières syndrome (AGS), which clinically mimics encephalopathies caused by *in utero*-acquired virus infections (11, 14, 28–33). Homozygous *Adar* gene knockout is embryonic lethal in mice, causing mass apoptosis of fetal liver hematopoietic cells by embryonic day 11.5-12.5 (20, 34). In contrast, *Adar^+/-^* mice are phenotypically normal and born at expected Mendelian ratios (18). Early fetal demise of *Adar ^-/-^* mice has been linked mechanistically to activation of the MDA5 pathway by dsRNA regions in cellular RNAs, chiefly repetitive elements, that are normally masked by A-I editing (15, 18, 20, 34, 35).

Indeed, constitutive MDA5 activation has been repeatedly observed to cause autoimmune syndromes such as AGS in both mice and humans (31, 36). In distinctive counterpoint, we have shown in a mouse model that chronic, systemic MDA5 activation caused by viral polymerase-generated dsRNA, which causes marked, lifelong ISG upregulation, can be well-tolerated, even when it is also strongly protective against viral diseases (37–39). The mice are transgenic for the RdRp of a neurovirulent mouse picornavirus (Theiler’s murine encephalomyelitis virus, TMEV) expressed under transcriptional control of the non-selective ubiquitin C promoter, which causes chronic dsRNA-triggered innate immune activation (37, 38). Tissues of the RdRp transgenic mice (RdRp^tg^ mice; see first Methods section for nomenclature) express low levels of the polymerase, which templates on host RNAs to synthesize dsRNA. Elevated dsRNA is detected in RdRp^tg^ mouse tissues using the K1 anti-dsRNA antibody (37). In addition, a catalytic center mutant of the TMEV RdRp lacked all ISG up-regulating activity (37). RdRp^tg^ mice have global, high upregulations of ISGs and robust protection against ordinarily lethal challenges by a variety of RNA and DNA viruses, including Theiler’s virus itself, EMCV, vesicular stomatitis virus, a DNA herpesvirus (pseudorabies virus), and Friend retrovirus (37, 38, 40, 41). The model is mouse strain-independent, with equivalent phenotypes in FVB/NJ, BALB/c and C57BL/6J mice.

RdRp^tg/-^ and RdRp^tg/tg^ mice develop normally, with onset of the major ISG expression profile shortly after birth (39). The intriguing lack of deleterious effects from their chronically elevated ISGome differs strikingly from other constitutive MDA5 activation states (36). Body size, morphology, organ histology and longevity are normal (37–39). Crosses with *Ifih1*^-/-^ mice showed that the sustained innate immune activation is strictly dependent on MDA5, which is congruent with MDA5 but not RIG-I being the sensor of picornaviral dsRNAs (1, 5–8). It is further dependent on the downstream adaptor MAVS, and the type I IFN receptor (IFNAR1) and is abolished by knockout of these genes (37). Although type I IFNs are not detectably over-expressed in RdRp^tg^ mice tissues, antibody-mediated blockade of the type I IFN receptor in adults terminates the ISG profile, indicating that some ongoing IFN-I signaling is required to sustain it (38, 39). *TLR3*, *IFNGR1* and *RAG1* knockout crosses also showed the ISG profile does not depend on TLR3, interferon gamma signaling or the adaptive immune system. As expected from the complete abrogation of ISG upregulation by *Ifih1* knockout, *Ddx58* knockout mouse crosses further confirmed no dependence on RIG-I (our unpublished data). Thus, the model is distinctive in that it represents a pure dsRNA-induced MDA5 hyperactivation state, and also one that is mediated through the wild type MDA5 sensor, and via “viral” dsRNA.

Prompted by the well-tolerated ISG elevations in this model, we here carried out experiments that demonstrate that RdRp^tg^ mice also resist induction of SLE in the BM12 lupus model, which we find is linked to increased quantities and effector function of regulatory T cells (T_reg_). We show that introducing single *Adar* allele knockout – which, similar to RdRp transgenesis, produces no abnormalities by itself – breaks the RdRp^tg^ protective state, yielding a severe autoinflammatory disease. RdRp^tg^ mice lacking one *Adar* allele have stunted growth, gray fur, abnormal dental and skeletal structures, failure to thrive, highly dysregulated ISG expression, and abnormal A-to-I editing. This dsRNA-driven, MDA5-wild type model establishes that viral polymerase-sourced dsRNA can drive interferonopathy pathogenesis and illuminates the autoimmunity preventing role of ADAR1.

## RESULTS

### RdRp^tg^ mice resist induction of SLE

Since picornaviral RdRp^tg^ mice do not exhibit apparent fitness “costs” of their chronic immune system activation (37–39), an unanswered question is whether they may be more autoimmunity-prone like other constitutive MDA5 activation models (36, 42, 43). To address this experimentally, we carried out a strong autoimmunity provocation by adopting the BM12 inducible SLE model, in which lupus is initiated by injection of splenocytes from MHC class II-mismatched mice (44, 45). Wild type (WT) and RdRp^tg/-^ mice were injected with 1×10^8^ splenocytes isolated from BM12 mice or were control-injected with PBS and evaluated for disease two weeks later. One of the clearest morphological changes in the BM12 model is splenomegaly, which results from germinal center expansion. Spleens were harvested and weighed before processing for flow cytometric analyses. As anticipated, we observed a significant increase in spleen size in WT mouse BM12 splenocyte recipients, indicative of lupus-like disease initiation (**Fig. 1A**). In contrast, RdRp^tg^ mouse BM12 splenocyte recipients had only minor, statistically non-significant increases in spleen size (**Fig. 1A**, p = 0.126 as compared with PBS-injected RdRp^tg^ mice).

**Figure 1.**
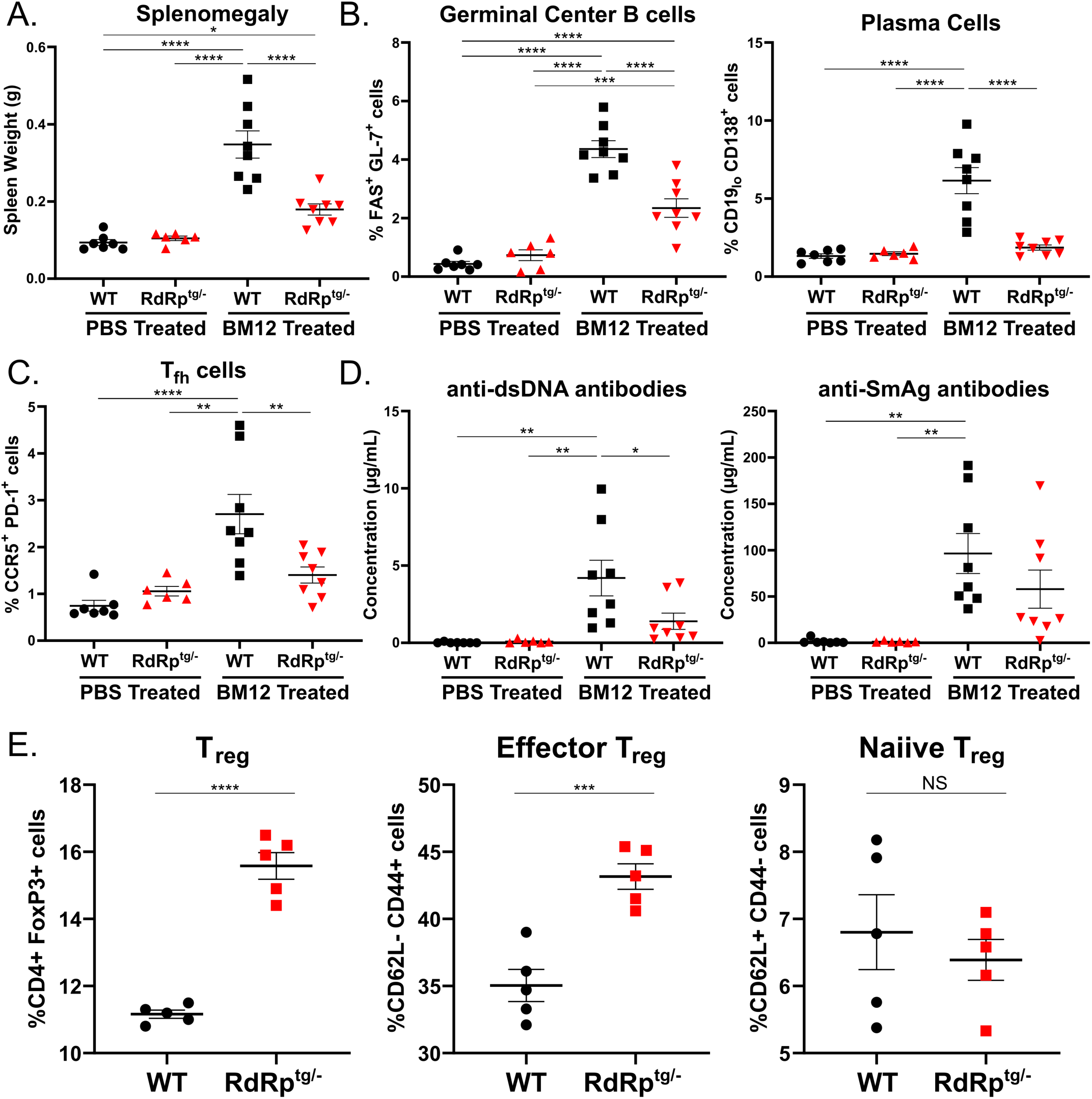
RdRp^tg^ mice resist SLE-like disease induction in the BM12 model of lupus. Ten-week-old mice were challenged with 100 million BM12-derived splenocytes. Mice were harvested 14 days post challenge for analyses. **(A)** Splenomegaly. Weights of BM12 and sham-injected mice spleens were measured. **(B,C)** Germinal center B cells, plasma cells and Tfh cells. RBC-lysed, single cell suspensions of splenocytes from BM12 or sham injected animals were generated for flow staining and germinal center B cells and plasma B cells (B), or follicular helper cell populations (C) were measured by flow cytometry according to gating described in materials and methods section. **(D)** Anti-nuclear antibodies. Sera from BM12 splenocyte-injected or sham injected mice were used to measure anti-dsDNA antibodies and anti-SmAg antibodies by quantitative ELISA. Data in A-D are from n = 7, 6, 8, 8 mice for WT (PBS), *RdRp^tg/-^*(PBS), WT (BM12), and *RdRp^tg/-^*(BM12) groups respectively. **(E)** T_reg_ cells. RBC-lysed single cell suspensions were generated from spleens of untreated, ten-week-old WT or RdRp^tg^ mice (n = 5 for each group) and T_reg_ subsets were determined by flow cytometry. Data were analyzed using one-way ANOVA followed by Tukey tests for (A-D) and an unpaired student’s T test for (E). * = p < 0.05, ** = p < 0.01, *** = p < 0.001, **** = p < 0.0001. Data points represent individual animals, graphs show means with standard deviations (s.d.).

We next examined cellular subsets in the spleen, specifically germinal center B cells and plasma B cells, which expand as they become activated to secrete anti-nuclear antibodies (ANAs), and T follicular helper (T_fh_) cells, which provide antigenic stimulus in germinal center reactions (46, 47). For all three cell populations, WT mouse BM12 splenocyte recipients developed significant increases, indicative of ongoing germinal center reactions and increased antibody production (**Fig. 1B,C**). In contrast, and paralleling the spleen measurements, RdRp^tg^ mouse BM12 splenocyte recipients developed little to no increases in all three cellular populations. Significant increases over sham-treated animals were only seen in germinal center B cells (**Fig. 1B,C**). Minor changes in overall B cell and T cell populations were also observed, but none that account for the drastic differences in cellular subsets seen in WT BM12 splenocyte recipient animals (**Fig. S1A,B**).

To corroborate the results, we collected serum from splenocyte-recipients and control PBS recipients to measure production of ANAs, which are a hallmark of SLE and drive pathology in this model (44, 45). We measured ANAs against double-stranded DNA (dsDNA) and Smith antigen (SmAg), both of which are frequently observed in SLE patients and SLE mouse models (47, 48) (**Fig. 1D**). Similar to the changes in cellular populations, there were significant increases in both of these autoantibodies in WT BM12 splenocyte recipients compared to the PBS recipient controls. In contrast, the RdRp^tg^ mouse BM12 splenocyte recipients produced less of both autoantibodies.

### T_reg_ cell expansion

Our previous investigations did not reveal significant differences in immune cell subsets between WT and RdRp^tg^ mice (38). Here we extended the studies to assess autoimmune suppressor cells, specifically regulatory T cells (T_regs_), which are critical for maintaining immune-tissue homeostasis. Knockout of T_regs_ results in disseminated autoimmune disease followed by rapid organismal decline and death (49, 50). We measured T_reg_ cell subsets in age-matched animals. There was a highly significant increase in T_reg_ cells in RdRp^tg/-^ mice compared to WT and specifically an increase in mature, effector T_regs_ (**Fig. 1E**). T_reg_ cell expansion in RdRp^tg^ mice may aid their tolerance to chronic innate immune activation.

The combined cellular and ANA data indicate that RdRp^tg^ mice are better able to control disease induction after an autoimmune provocation than WT mice. This result is distinctive compared to other MDA5-pathway driven hyperimmune mice models, which have been shown to be equally or more susceptible to autoimmunity induction (42, 43)

### Investigation of ADAR1

In addition to Treg cells, we suspected that mechanisms of protection at the intrinsic cellular level are likely to provide key regulation. Given its known role in preventing activation of the MDA5 pathway by double stranded segments of endogenously encoded RNAs, we hypothesized that ADAR1 may prevent adverse inflammatory sequelae in RdRp^tg^ mice. First, we determined whether *Adar* expression is affected by RdRp genotype status. *Adar* encodes two functional isoforms, ADAR1 p150, which edits transcripts in the cytoplasm and is IFN-inducible, and constitutively expressed ADAR1 p110, which is generated by alternative splicing or by translation from the p150 mRNA by leaky internal ribosome scanning (51) and edits transcripts in the nucleus (19, 52, 53). We observed no significant difference at either the mRNA or protein levels in expression of the p110 isoform in RdRp^tg^ mice as compared to WT mice (**Fig. 2A,C**). In contrast, there was a marked increase in expression of the IFN-inducible p150 isoform (**Fig. 2B,C**). These results suggested that ADAR1 p150, itself an ISG, is upregulated in RdRp mice, perhaps helping to modulate the ISG activation in these animals.

**Figure 2.**
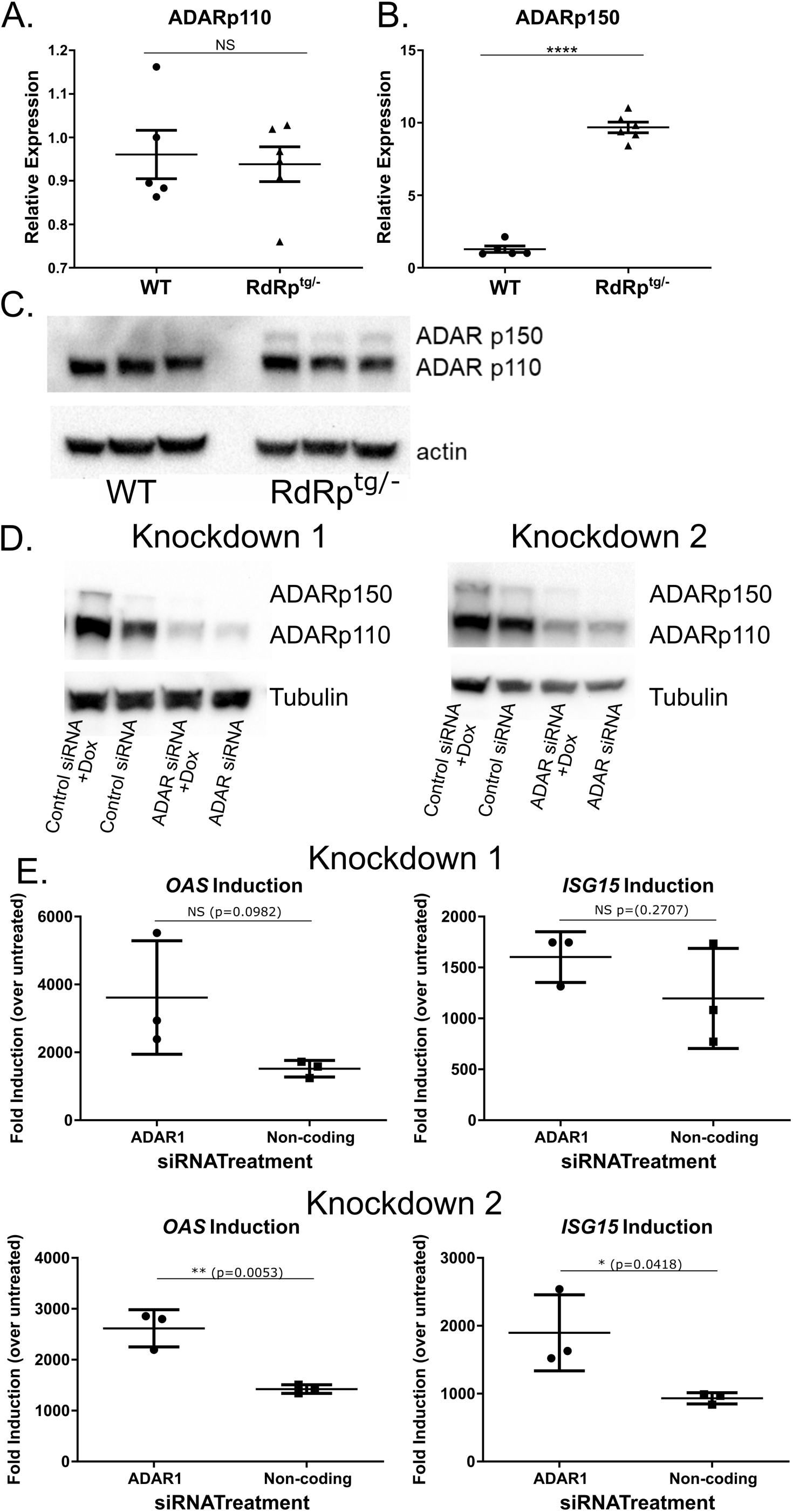
ADAR p150 and ADAR p110 levels in RdRp^tg^ mice, and effects of *ADAR* knockdown on RdRp-dependent ISG expression. RNA isolated from brains of 4 to 5 week old WT or *RdRp^tg/-^* mice (n = 5 for WT, 6 for *RdRp^tg/-^*) was used to measure **(A)** ADAR p110 and **(B)** ADAR p150 transcripts by qPCR. Data points represent individual animals, graphs shown means, s.d. **(C)** Immunoblotting for ADAR p110 and ADAR p150 in 4 to 5 week old mouse WT and *RdRp^tg/-^*brain. n = 3 animals per genotype. **(D)** A549 cells with inducible Theiler’s virus RdRp (Tet-on system). Data are shown from two representative knockdowns. **(E)** RNAs isolated from the parallel knockdowns done in panel D were used for qPCR analysis to determine relative levels of the ISGs *OAS* and *ISG15* mRNAs. mRNAs were harvested 54 hours after *ADAR*-targeting siRNA addition and 48 hours after dox addition. Comparisons were made between dox-treated and dox-untreated cells and expressed as fold mRNA changes induced by dox. Transfection controls (cells receiving transfection reagents but no siRNA) showed similar levels of ISG induction after dox treatment as control siRNA-transfected cells, indicating lack of contribution of transfected siRNAs to immune activation. Data are means and s.d. of triplicate biological replicates with three technical replicates each. NS: not significant; * = p < 0.05, ** = p < 0.01, **** = p < 0.0001; unpaired Student’s T test.

We next knocked down *ADAR* expression in human lung epithelial cells (A549) in which we have engineered and validated inducible TMEV RdRp expression under the control of a doxycycline (dox) inducible promoter (37). These cells broadly upregulate ISGs in response to RdRp induction in a pattern very similar to the RdRp^tg^ mouse (37). In two separate experiments, we achieved knockdowns of ADAR p110 and p150 isoforms (**Fig. 2D**). Dox induction of the RdRp after siRNA transfection led to over three log_10_ increases in mRNAs for two classical ISGs, *OAS* and *ISG15*, which were moderately accentuated by *ADAR* depletion in these short-term experiments (**Fig. 2E**). We therefore proceeded to generate mice with *Adar* gene knockouts.

### *Adar* haploinsufficiency, by itself well-tolerated, causes multi-system disease in RdRp^tg^ mice

As noted, by two weeks after birth, RdRp^tg^ mice develop high, sustained upregulations of many ISGs and subsequently tolerate them throughout their (normal) lifespans (37, 39). To determine whether ADAR1 is involved in the protection against deleterious effects of autoinflammation, *RdRp*^tg/tg^ mice were crossed with *Adar*^+/-^ mice. *RdRp^tg/-^* mice also have elevated ISGs, albeit with more variability in fold-induction than RdRp^tg/tg^ mice (for example, see **Fig. 2B,C** for expression of the ISG ADAR1 p150, and Painter et al. (37)). Since in our hands and others *Adar*^+/-^ mice have no phenotypic abnormalities, we did not anticipate a major effect in first-generation crosses even in the presence of an RdRp-mediated ISG response, since the F1 animals will retain a functional *Adar* allele.

However, that was not the outcome. First, there were major gross phenotypic differences between *RdRp^tg/-^ Adar^+/-^* mice and littermate *RdRp^tg/-^ Adar^+/+^* controls (**Fig. 3A-C**). The RdRp^tg^ mice lacking one *Adar* allele were significantly smaller in size (**Fig. 3A, B**), with gray instead of black fur (**Fig. 3A, C**). These features were equivalent in males and females (**Fig. 3A, C**). *RdRp^tg/-^ Adar^+/-^* mice also have small, poorly developed, misshapen teeth (**Fig. 3C**). Runting was unrelated to the dental abnormalities, since suckling behavior was unaffected. In addition, the size evaluations were done at the time of weaning, and post-weaning mice were kept on a wet chow diet, which they were also observed to ingest equivalently. *Adar^+/-^* mice were normal in appearance as expected.

**Figure 3.**
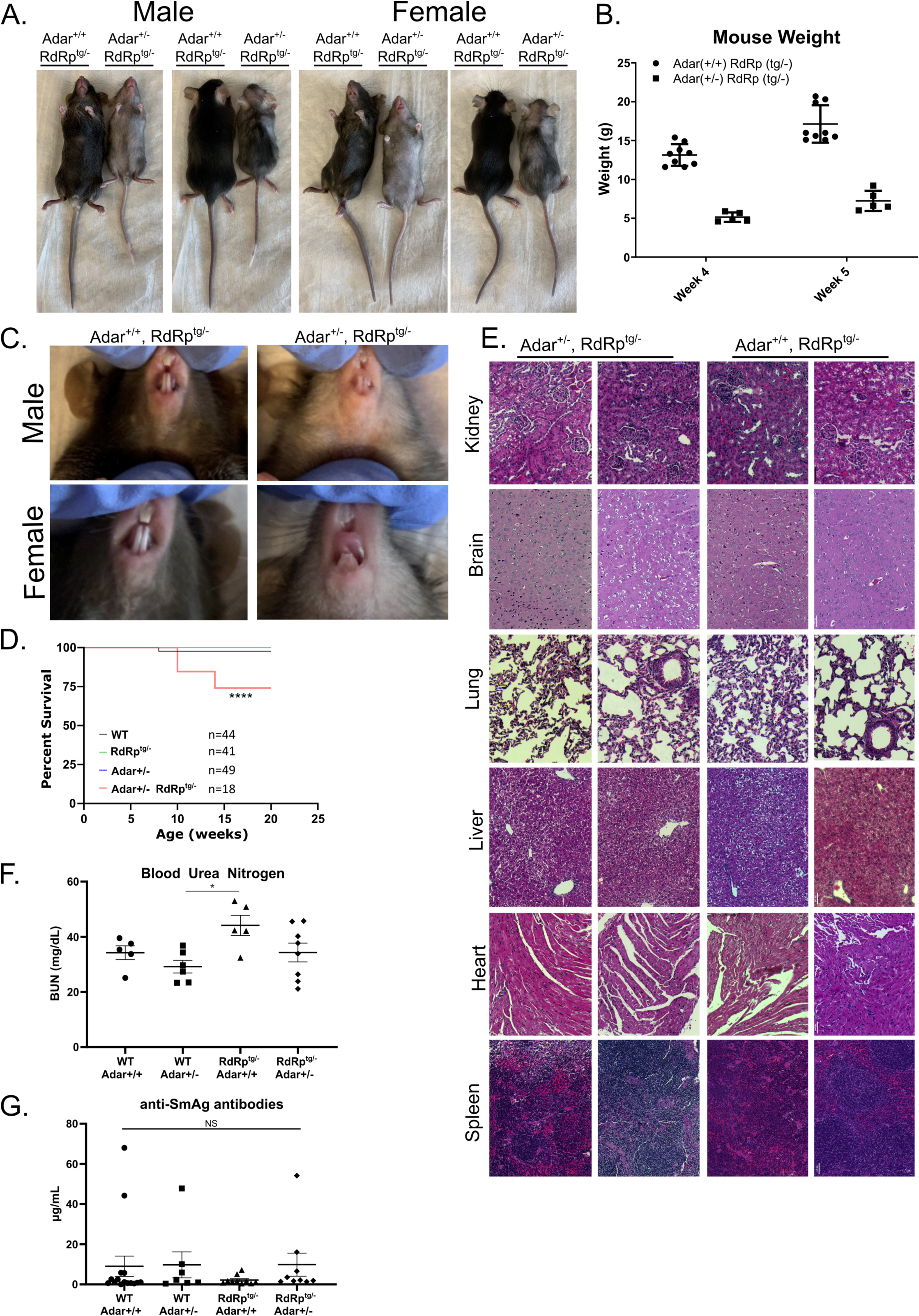
Phenotypic and histological differences in RdRp^tg/-^ Adar^+/-^ mice. **(A)** Size and coat color differences in 5-week-old, littermate *RdRp^tg/-^*and *RdRp^tg/-^ Adar^+/-^* mice. **(B)** Weight differences between 4 week and 5-week-old littermate *RdRp^tg/-^* and *RdRp^tg/-^ Adar^+/-^* mice (n = 9 and 5 respectively). **(C)** Dental developmental differences between *RdRp^tg/-^*and *RdRp^tg/-^ Adar^+/-^* mice. **(D)** Survival of animals from each group, followed for 20 weeks and shown as Kaplan-Meier plot. **(E)** H & E-stained tissue sections from kidney, brain, lung, liver, heart and spleens from *RdRp^tg/-^* and *RdRp^tg/-^ Adar^+/-^* littermate mice. Pathological grading revealed no significant scoring in *RdRp^tg/-^Adar^+/-^* animals. **(F)** Blood Urea nitrogen from serum from 4-5 week old animals in WT, *Adar^+/-^*, *RdRp^tg/-^*, *Adar^+/-^RdRp^tg/-^* mice. n = 5, 6, 5, 8 for WT, *Adar^+/-^*, *RdRp^tg/-^*, *Adar^+/-^ RdRp^tg/-^*, respectively. **(G)** Anti-SmAG antibodies from serum from 4-5 week old animals as measured by quantitative ELISA. Four outlier high values were seen, but two were in the WT group. n = 15, 7, 10, 9 for WT, *Adar^+/-^*, *RdRp^tg/-^*, *Adar^+/-^ RdRp^tg/-^*, respectively. Anti-dsDNA antibody levels from matched serum was below the limit of detection for all animals tested. Where not indicated, all data and tissue sections come from a mix of male and female mice. Data in (B) were analyzed by two-way ANOVA where ****p < 0.0001. Data in D were analyzed using a log-rank test (Mantel-Cox) where **** = p < 0.0001. Data in (F) and (G) were analyzed by one-way ANOVA where *p < 0.05; data points represent individual animals, graphs show means and s.d.

Because of these phenotypic differences, we carried out tissue dissections. Survival of the *RdRp^tg/-^Adar^+/-^* mice began to fall significantly by 10 weeks after birth (**Fig. 3D**), so these studies were done at 4-5 weeks. Soft viscera including heart, lungs, liver, spleen, intestines, kidney and brain were proportionately small but the organ morphologies were otherwise not different from WT mice. Histopathology analysis of tissue sections by a veterinary pathologist did not reveal evidence for tissue inflammation or other abnormalities (**Fig. 3E**; see Supplemental Methods for scoring and data). Since some MDA5-driven mouse autoimmune models manifest with kidney impairment, blood urea nitrogen (BUN) values were determined at 4-5 weeks of age. Mean BUN was slightly higher in RdRp^tg^ mice but was not in either *Adar^+/-^*group (**Fig. 3F**). We also measured serum ANAs and found no significant increases in anti-dsDNA antibodies (data not shown as levels for all animals were below the limit of assay detection) or anti-SmAg antibodies (**Fig. 3G**). In addition to the abnormal teeth, necropsy revealed thin, pale long bones suggesting decreased bone density. Bone frailty was also readily apparent during cervical dislocations, which subjectively required much less force Similar dental and bone abnormalities can occur in Singleton-Merten Syndrome (SMS), a rare human autosomal dominant innate immune disorder consisting of musculoskeletal abnormalities (osteopenia, osteoporosis, skull thickening, small stature, ligament frailty), dental anomalies (poor, primarily anterior teeth formation), variable arterial calcification, inflammatory skin changes (psoriasis), and a characteristic facies (high anterior hair line, broadened forehead, asymmetric ptosis, smooth philtrum, and thin upper vermillion) (54, 55). Genetic studies of individuals with SMS have revealed causation by a subset of single amino acid missense gain-of-function (GOF) mutations in MDA5 (R822Q, A489T, T331I/R), which cause constitutive activation (56–60). There is inter-familial and intra-familial variation in syndromic manifestations, with partial penetrance evident in some kindreds(55). and overlap with Aicardi-Goutières syndrome has been reported (58, 59, 61). Similarly, two families with atypical SMS lacking the dental abnormalities were found to have constitutively active single amino acid mutants of RIG-I rather than MDA5 (62).

As *RdRp^tg/-^ Adar^+/-^* mice had micrognathia as well as small, misshapen and partially translucent incisors (**Fig. 3C**), µCT imaging and mechanical property assessments of femurs were performed to determine whether *RdRp^tg/-^ Adar^+/-^* mice also had altered underlying changes to bone architecture or properties. Consistent with the runting observed in 5-week-old mice (**Fig. 3A,B**), femurs of 6-week-old *RdRp^tg/-^ Adar^+/-^* were significantly shorter than control groups (**Fig. 4A,B**). As in other RLR-related genetic disorders (55, 62), weight, femur length, and other mouse characteristics were variably penetrant. Bartlett’s test demonstrated a statistically significant (p=0.0142) difference in homoscedasticity for femur length, yet a Kolmogorov-Smirnov test did not identify significant (alpha=0.05) deviation from normal distribution for *RdRp^tg/-^ Adar^+/-^* or other groups. Thus, we did not separate *RdRp^tg/-^ Adar^+/-^* mice into ‘penetrant’ and ‘non-penetrant’ subsets in subsequent analyses and graphics, but we labeled mice with gray fur with light blue symbols in Fig. 4. *RdRp^tg/-^ Adar^+/-^* femurs had significantly reduced cortical bone volume fraction and correspondingly reduced measures of cortical thickness, perimeter, area and porosity (**Fig. 4C**). In contrast, trabecular number and trabecular separation were not different between groups (**Fig. 4D-E**). Mechanical properties of the femurs were further investigated using three-point bending to failure. Stiffness and maximum load of *RdRp^tg/-^ Adar^+/-^*femurs were significantly reduced compared to control groups (**Fig. 4F,G**). Modulus and ultimate stress of the femurs were calculated from mechanical properties and mid-diaphysis cross-sectional geometry from µCT. In contrast to the reduced mechanical properties observed, these material properties of *RdRp^tg/-^ Adar^+/-^* femurs were not altered (**Fig. 4I-J**), nor was the total cortical mineral density calculated from the µCT imaging (**Fig. 4H**). These results indicate that double heterozygosity (*RdRp^tg/-^ Adar^+/-^*) led to femurs having inferior stiffness and strength stemming from smaller size but not from impaired bone material quality, in addition to their dental maldevelopment and fur graying.

**Figure 4.**
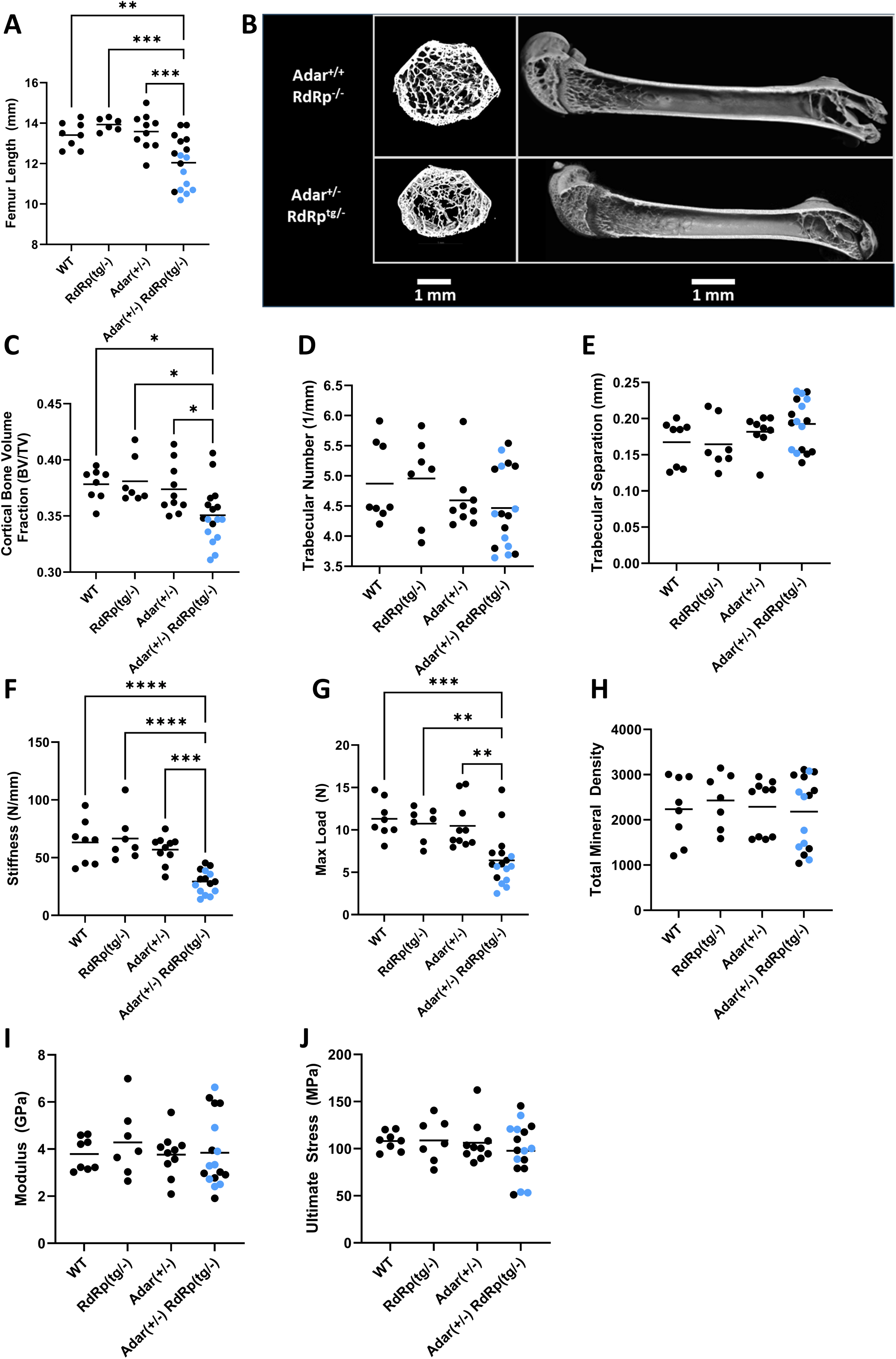
Skeletal and dental features of wild type, *RdRp^tg/-^*, *Adar^+/-^*, and *Adar^+/-^ RdRp^tg/-^* mice. Femurs from 6-week-old animals were manually de-fleshed and used for all analyses. **(A)** Femur lengths of WT, *Adar^+/-^*, *RdRp^tg/-^,* and *RdRp^tg/-^ Adar^+/-^* mice (n = 8, 6, 10, and 17 animals respectively). **(B)** Representative µCT images, distal femur sections and whole bone. **(C)** Cortical bone volume fraction (BV/TV), n = 8, 7, 10, 17. **(D)** Trabecular number, n = 8, 7, 9, 17. **(E)** Trabecular separation, n = 8, 7, 9, 17. **(F)** Stiffness, n = 8, 7, 10, 15. **(G)** Maximum load, n = 8, 7, 10, 17. **(H)** Cortical total mineral density, n = 8, 7, 10, 17. **(I)** Modulus, n = 8, 7, 10, 17. **(J)** Ultimate stress, n = 8, 7, 10, 17. Mice with gray fur are indicated by light-blue symbols. Data were analyzed first by using a ROUT test to remove outliers (Q = 1%), which resulted in removal of two mice from one group in one panel (the *RdRp^tg/-^ Adar^+/-^* group in Fig. 4F, stiffness testing; hence there are 15 mice as opposed to the 17 for this genotype in the other panels). A one-way ANOVA comparing *RdRp^tg/-^ Adar^+/-^* to each other group was used followed by a Tukey tests where * = p < 0.05, ** = p < 0.01, *** = p < 0.001, **** = p < 0.0001. Data points represent individual animals, horizontal bars indicate means.

Resemblance to classical human SMS was partial since we the mice did not develop aortic or cardiac valvular calcification, a core feature of the human syndrome, nor was there aberrant brain calcification (**Fig. S2**) as occurs in AGS (11) and in some SMS-AGS overlap cases(58, 59, 61). Psoriasis is less commonly observed in SMS patients and was not observed in *RdRp^tg/-^ Adar^+/-^*animals, leaving fur graying as the identifiable integument abnormality. Glaucoma can also occur in SMS, but intraocular pressures measured by tonometry in affected mice were normal and not significantly different from WT mice (data not shown).

### Transcriptional changes in *RdRp^tg/-^ Adar^+/-^* mice are consistent with an interferonopathy

The SMS-resembling phenotype triggered by the combination of one RdRp transgene allele and one *Adar* null allele, with neither alone producing abnormalities, prompted us to examine tissues further for differences that may be driving pathogenesis. We performed RNA-seq on brain tissue, with four biological replicates from each of the four genotypes WT, *Adar^+/-^*, *RdRp^tg/-^* and *RdRp^tg/-^ Adar^+/-^*. Brains were analyzed because we previously observed that brains of RdRp^tg^ mice have particularly high ISG elevations compared to other organs, although gene expression changes are qualitatively similar and substantial across all major organs (39). Here, in multidimensional scaling plots, we observed that the samples cluster strongly by genotype along the M1 axis, which validates that animals of the same genotype have similar gene expression profiles (**Fig. 5A**). The double heterozygotes (*RdRp^tg/-^ Adar^+/-^*) cluster the farthest from any other genotype, suggesting they have distinctly different RNA expression patterns. Assessing expression changes across the groups, the greatest differences were between *RdRp^tg/-^ Adar^+/-^* mice relative to WT mice and relative to *Adar^+/-^*mice, with the majority of upregulated genes being known ISGs (136 out of 151 and 124 out of 137, respectively) (**Fig. 5B,C, Fig. S3**). These outcomes suggest that *RdRp^tg/-^ Adar^+/-^* mice have an extreme ISG response that likely leads to the Singleton-Merten interferonopathy-resembling phenotype. Interestingly, when we examined which genes are changing across genotype comparisons, we saw a largely overlapping set of ISGs upregulated in the presence of the RdRp transgene or in the absence of one *Adar* allele (**Fig. 5D**). All three non-WT genotypes (*Adar^+/-^, RdRp^tg/-^,* and *RdRp^tg/-^ Adar^+/-^*) showed significant upregulation of overlapping ISGs, but the magnitude of upregulation varies drastically by genotype (**Fig. 5D**). Consistently the dual heterozygotes have extremely high induction, elevated 3-5-fold over what we observed in *RdRp^tg/-^* mice, although largely concordant in which genes are induced, while *Adar^+/-^* have only low level ISG induction (**Fig. 5B,D**). Thus, synergy between the RdRp transgene and the absence of one *Adar* allele leads to a hyper-induced ISG state in mice that is no longer tolerated. We considered whether differing expression of the TMEV RdRp transcript itself might be influencing the observed double heterozygote differences (even though the ubiquitin C promoter controls its transcription), but the RdRp mRNA was similarly elevated in both (**Fig. 5E**).

**Figure 5.**
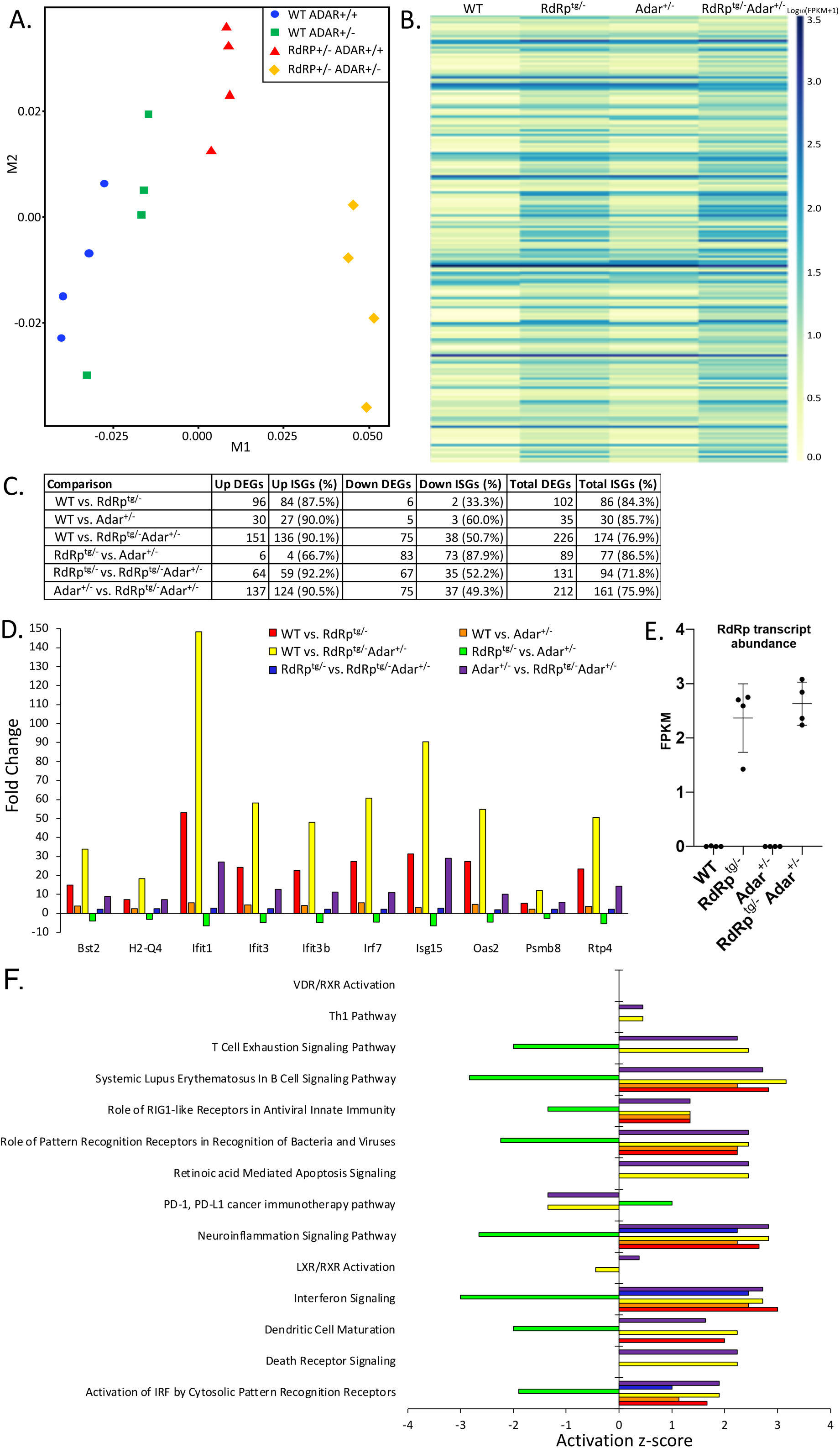
RNA-seq analysis reveals clear pattern of ISG upregulation suggestive of an interferonopathy. RNA-seq from neuronal tissue from n = 4 age-matched (5 weeks) mice per group. Where possible, littermate controls were used for analysis. Two male and two female mice were used per group. **(A)** Multi-dimensional scaling plot of RNA expression profiles in WT-WT, *Adar^+/-^*, *RdRp^tg/-^,* and *RdRp^tg/-^ Adar^+/-^* mice. **(B)** Heatmap of differentially expressed ISGs across the four genotypes (**Fig. S3** shows an enlarged version with gene names annotated). FPKM values were log transformed with 1 pseudocount to facilitate visualization. **(C)** Summary table of differentially expressed genes (DEGs) and proportions of which are known ISGs as determined by the Interferome database. **(D)** Differentially expressed genes that are shared across all group comparisons. **(E)** RdRp mRNA abundance in the four genotypes. **(F)** Canonical molecular pathways from IPA that are significant across group comparisons.

### Pathway Analysis

Ingenuity Pathway Analysis (IPA) was used to assess which canonical molecular pathways are activated or inhibited. *Adar^+/-^, RdRp^tg/-^,* and *RdRp^tg/-^ Adar^+/-^* mice showed activation of genes related to interferon signaling, neuroinflammatory signaling and PRRs (**Fig. 5F**). Genes associated with dendritic cell (DC) maturation were activated in *RdRp^tg/-^* and *RdRp^tg/-^ Adar^+/-^*mice (**Fig. 5F**). Increased DC maturation, specifically in *RdRp^tg/-^ Adar^+/-^*animals, may contribute to the observed disease. We also carried out analyses of upstream regulators that may be responsible for gene expression patterns. In WT-*Adar^+/^*^-^, *RdRp^tg/-^ Adar^+/+^,* and *RdRp^tg/-^ Adar^+/-^* mice, we not surprisingly identified *Ddx58* (RIG-I), and *Ifih1* (MDA5) as upstream regulators activated in those samples (**Fig. S4**). Main intermediates in interferon signaling pathways, including IRF7, IRF9, STAT1, and STAT2 were also statistically significant predicted key regulators. (**Fig. S4**). Only in mice missing one *Adar* allele (*Adar^+/-^* and *RdRp^tg/-^ Adar^+/-^* mice) did we also identify predicted regulation by additional PRR proteins, including TLR3, which recognizes dsRNA, and *ZBP1* (DAI), which recognizes Z-form nucleic acids (63), both resulting in activation of interferon signaling (**Fig. S4**). Interactions with other nucleic acid sensing molecules besides MDA5 may thus play roles in sensing of ADAR1-edited transcripts.

### Molecular and cellular differences in *RdRp^tg/-^ Adar^+/-^* mice validate RNA-seq analyses and are consistent with interferonopathy

To characterize the RNA-seq results further, we determined levels of ISG mRNAs in brain tissues of *Adar*^+/+^ and *Adar*^+/-^ mice, with or without the RdRp transgene. Similar to the RNA-seq data, in *Adar*^+/-^ mice, there were only small increases in levels of three classical ISG mRNAs compared to WT mice (about 2- to 5-fold; **Fig. 6A**). In contrast, and consistent with prior data (37–39). *RdRp^tg/-^* mice had major increases compared to WT mice. However, the dual heterozygotes (*RdRp^tg/-^ Adar^+/-^*) had much higher ISG mRNA elevations, on the order of 200 – 600 fold relative to WT.

**Figure 6.**
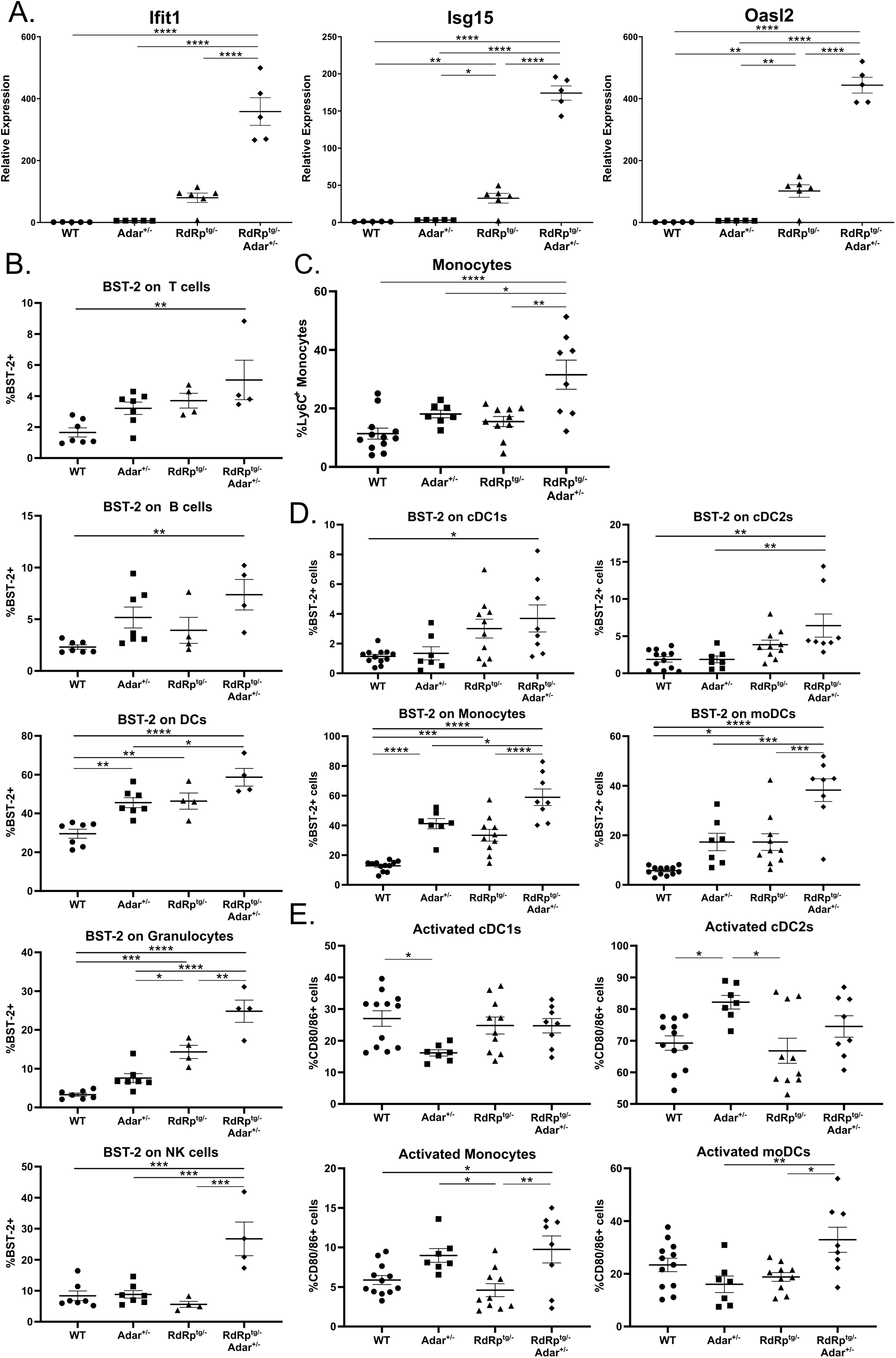
ISG and leukocyte subset differences in RdRp^tg/-^ Adar^+/-^ mice. **(A)** qPCR of three representative ISGs (*Ifit1*, *Isg15*, and *Oasl2*) in brain tissue of 4-5 week old WT, *Adar^+/-^*, *RdRp^tg/^*, and *RdRp^tg/-^ Adar^+/-^* mice (n = 5, 5, 6, 5, respectively). **(B-E)** Flow cytometry analysis of cellular populations derived from the spleens of 4-5 week old mice. Single cell, RBC-lysed solutions were prepared for use in analysis. **(B)** BST-2 expression on the main immune cell subsets in the spleen including T cells (CD3^+^), B cells (CD19^+^), DCs (CD11c^+^), granulocytes (Ly6G^+^), and NK cells (NKp46^+^). n = 7, 7, 4, 4 for WT, *Adar^+/-^*, *RdRp^tg/^*, *RdRp^tg/-^ Adar^+/-^*, respectively. **(C)** Monocytes populations across all four groups. **(D)** BST-2 expression on monocyte/DC cell subsets including monocytes, cDC1s, cDC2s, and moDCs. **(D)** Activation (CD80/86 expression) of monocyte/DC subsets including monocytes, cDC1s, cDC2s, and moDCs. C-E: n = 12, 7, 10, 8 for WT, *Adar^+/-^*, *RdRp^tg/^*, *RdRp^tg/-^ Adar^+/-^*, respectively. For all graphs shown, data were analyzed using a one-way ANOVA followed by a Tukey tests to determine significance where * = p < 0.05, ** = p < 0.01, *** = p < 0.001, **** = p < 0.0001. Data points represent individual animals with the mean and s.d. shown as bars.

Importantly too, breeding to *Ifih1*^-/-^ mice demonstrated that loss of MDA5 resulted in rescue of animal growth and a complete abolishment of ISG upregulation observed in *RdRp^tg/-^ Adar^+/-^* animals (**Fig. S5**), reinforcing the MDA5-dependence of the phenotypes. Given the massively amplified ISG transcriptional pattern in the double heterozygotes, and the immune cell activation and maturation pathways enrichment observed (**Fig. 5, Fig. S6**), we next harvested the spleens of 4-5 week old mice for flow cytometry analyses.

Aberrantly activated immune cell subsets are canonical autoimmune disease features and identifying them would support the conclusion of a dysregulated immune state in *RdRp^tg/-^ Adar^+/-^* mice. In major immune cell populations in the spleens, including B cells, T cells, neutrophils, DCs, and granulocytes, we found no major differences in cell population proportions across different genotypes (**Fig. S7A**). The antiviral protein BST-2 (Tetherin) is a classical ISG with a high dynamic range of expression in response to type I interferon stimulation in multiple mammals, including mice (64–66), and the mRNA was consistently upregulated across comparisons in the RNA-seq experiments. We therefore determined cell surface BST-2 protein expression in the various immune cell populations, and found it to be elevated on all cell types of *RdRp^tg/-^ Adar^+/-^* mice and was slightly upregulated on DCs in WT-*Adar^+/-^*and *RdRp^tg/-^* animals and granulocytes from *RdRp^tg/-^*animals (**Fig. 6B**), similar to what we have observed previously (38). We also examined monocyte and dendritic cell differentiation, maturation and activation in these animals (**Fig. 6C, D, E**). Monocytes were significantly increased in *RdRp^t/-^ Adar^+/-^* animals (**Fig. 6C**) and cDC1s were slightly decreased also (**Fig. S7B**). Monocytes are circulating precursor cells that infiltrate into the tissue upon detection of proinflammatory stimuli and differentiate into effector monocyte-derived DCs (moDCs). An increase in this cell population is suggestive of increased demand for precursor cells in these animals. In support of this hypothesis, we observed increased BST-2 expression on all DC subsets from *RdRp^tg/-^ Adar^+/-^* mice, as well as increased expression of the activation markers CD80/86 on *RdRp^tg/-^ Adar^+/-^*monocytes and moDCs (**Fig. 6D, E**). Disorders in DC function and activity have been widely implicated in several autoimmune diseases, including SLE, and may contribute to disease development in *Adar*^+/-^ RdRp^tg/-^ animals.

The immune cell profiling above and the upregulated ISG profiles raised the question of whether proinflammatory cytokines are elevated in the double heterozygotes. Indeed, transcripts for IFNβ, which were undetectable at baseline in WT and Adar+/- mice, were detectably elevated, but at low levels, in both RdRp^tg/-^ and more so in *RdRp^tg/-^ Adar^+/-^* mice (**Fig. S8**). While classical proinflammatory cytokines such as IL6 and TNFα were not elevated in any group, several proinflammatory C-C and C-X-C chemokines were, most prominently CXCL10, CXCL11, CCL2 and CCL5 (**Fig. S8**).

### Editome analyses show dual heterozygotes have dysregulated A-I editing

To interrogate the relationship of the loss of one *Adar* allele to the SMS-like phenotype of *RdRp^tg/-^ Adar^+/-^* mice, we characterized genome-wide A-to-I editing differences between all four genotypes of animals (**Fig. 7**). We first carried out whole exome sequencing to eliminate any SNPs that may be unique to our colony or genotypes (of which very few were detected), as compared to the reference genome. We then identified all A-to-G changes in RNA-seq data (inosine is decoded as guanine during sequencing). We found that the majority of edited sites in all four genotypes were in intergenic regions and 3’UTRs, with a fractionally larger portion for 3’UTRs in RdRp^tg^ mice. Exons and introns were represented approximately equivalently (**Fig. 7A, B**). Interestingly and unexpectedly, while edited site locations were similarly proportioned among the four genotypes (**Fig. 7A**), the number of edited sites increased substantially (50 – 65%) in *RdRp^tg/-^ Adar^+/-^* mice compared to the other three genotypes (**Fig. 7B**). Numbers of edited genes also increased. This seemingly paradoxical result in the Singleton-Merten-affected dual heterozygotes was apparent across all gene regions. In contrast, in the *Adar^+/-^* and *RdRp^tg/-^*mice, the number of edited genes decreased compared to WT mice and the overall number of edited sites remained largely unchanged (**Fig. 7B**). Despite the number of edited sites being unchanged for WT, *Adar^+/-^*, and *RdRp^tg/-^* mice, there was a shift from intergenic to extragenic regions in these animals, and there was a substantial proportionate rise in 3’UTR editing in RdRp^tg/-^ mice of both genotypes, which was quantitatively greater in *RdRp^tg/-^ Adar^+/-^* mice.

**Figure 7.**
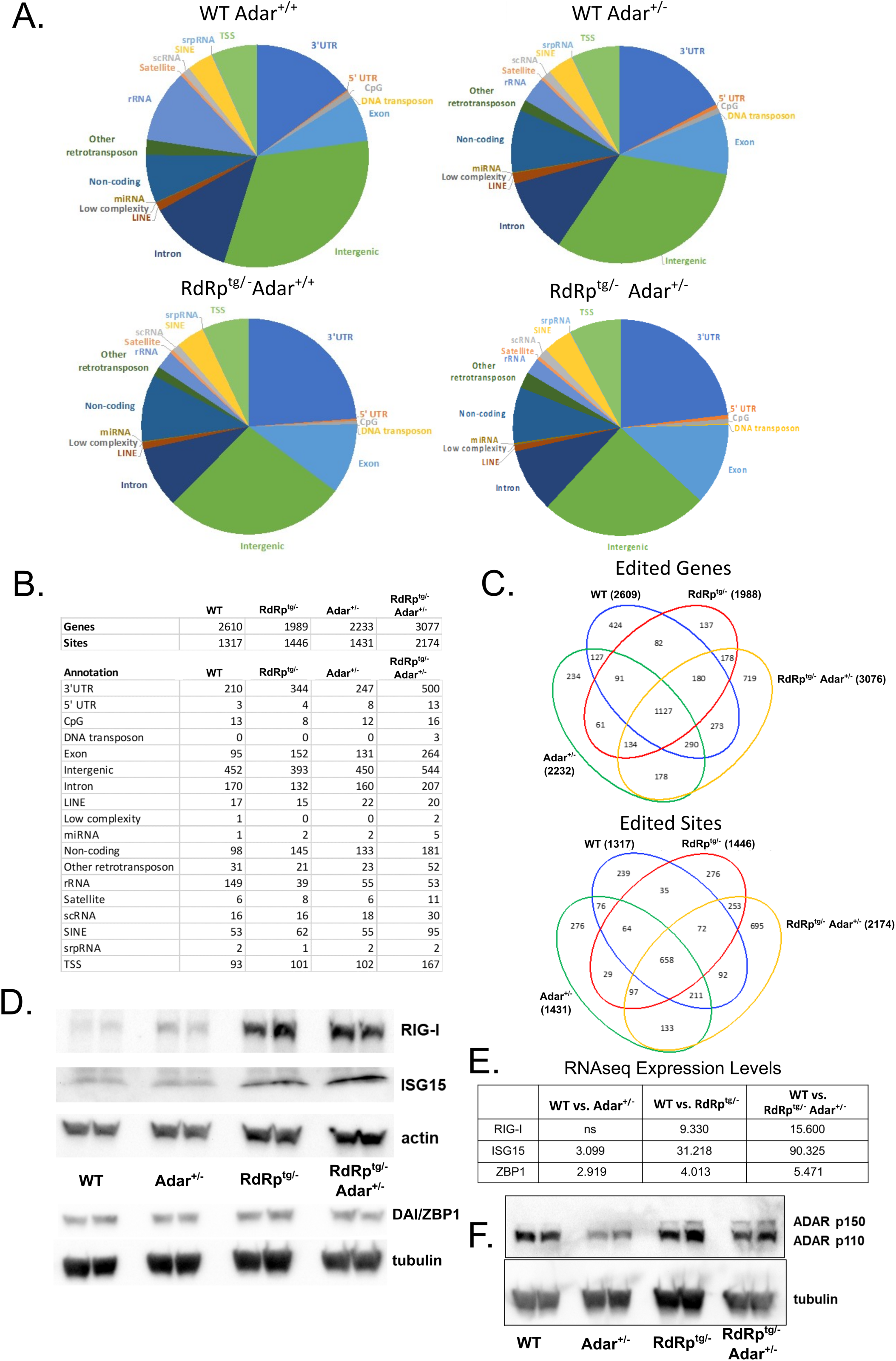
Analysis of the A-to-I editome reveals increased editing in *RdRp^tg/-^ Adar^+/-^* mice. Changes in A-to-I edits were determined via comparison of RNA-seq data with whole exome DNA sequencing. **(A)** Pie charts showing the proportion of editing sites within a given RNA element for each mouse genotype. **(B)** Table summary of the distribution of overall numbers of edited genes and sites for each genotype and the breakdown of locations of the edit sites among different RNA elements. For each of the four genotypes, four animals were sequenced and to be counted a gene must have been edited in all four animals sequenced but editing can occur anywhere in the gene. For sites, identical sites must be edited in all four animals of a genotype. **(C)** Venn diagrams showing overlap of edited genes (top) or sites (bottom) among the four genotypes. **(D,E)** Immunoblot analysis from two animals per group (D) and gene expression fold change (E) of three *Adar^+/-^ RdRp^tg/-^* uniquely edited genes in the RNA-seq data set (as compared to WT) in 5-week-old neuronal tissue. Rig-I, Isg15, and Zbp1 were evaluated as they are each ISGs that are highly upregulated in mice expressing RdRp and are key regulators of the antiviral response. **(F)** Expression of *Adar* isoforms in neuronal tissue of 4-5 week old mice from all four genotypes, as determined by western blot (n = 2 per group).

Because they possess only one functional *Adar* allele, it was unexpected that *RdRp^tg/-^ Adar^+/-^* mice would have these increases in the number of A-to-I edited genes and A-to-I edited sites (**Fig. 7B**). Thus, there is a dysregulation of A-to-I editing. When we examined overlap of edited sites or edited genes between genotypes (visualized in Venn diagrams, **Fig. 7C**), we observed the highest number of uniquely edited genes (719) and sites (695) in *RdRp^tg/-^ Adar^+/-^*mice, indicating that these animals have significantly different editing patterns compared to the other genotypes. It is notable that *RdRp^tg/-^ Adar^+/-^* mice have increased editing in LINEs, SINEs, DNA transposons and other retrotransposons, since increases in retroelement transcription are associated with autoimmune disease states (67). In addition, previous studies have shown ADAR1 to be critical for regulation of retroelement transcript self-reactivity(9, 10). Increased ADAR-specific editing of retroelements is normally protective, but in our double heterozygote mice this alone is not compensatory enough to prevent disease, indicating that the A-to-I editing process in these animals is abnormal.

Next, we used RNA-seq to analyze the changes in edited genes, along with relative expression in each group of mice, to determine associations with activated/inhibited canonical pathways and enriched diseases and biological functions (**Fig. S9**). Of edited mRNAs in *RdRp^tg/-^ Adar^+/-^* mice, genes associated with disease and biological functions related to apoptosis, cell death, and necrosis were enriched (**Fig. S9**). Notably, there was a clear reduction in genes involved in inflammatory disease states that was unique to comparisons with this *RdRp^tg/-^ Adar^+/-^* mice (**Fig. S9**).

To explore the question further, we asked how A-to-I editing of ISG mRNAs, when it occurs, affects expression of their encoded proteins in *RdRp^tg/-^ ADAR^+/-^* mice. We measured mRNA levels and performed protein immunoblotting for several ISGs that are uniquely edited in *RdRp^tg/-^ Adar^+/-^* mice: Rig-I, Isg15, and Zbp1. Both Rig-I (*Ddx58*) and Zbp1 (*Zbp1*) were also predicted by the RNA-seq analysis to be key upstream gene regulators (genes with known activation/inhibition effects on the enriched pathways; **Fig. S4**). Expression levels of Rig-I, Isg15, and to a much lesser extent and not at the protein level, Zbp1, were increased in *RdRp^tg/-^*and *RdRp^tg/-^ Adar^+/-^* mice, but not in *Adar^+/-^*mice (**Fig. 7D,E)** indicating that A-to-I editing does not reduce expression of these ISGs. Additionally, ADAR1 isoforms were expressed in *RdRp^tg/-^ Adar^+/-^* mice at relatively similar levels as their *wild type* counterpart (**Fig. 7E,F**), with slightly higher p150 protein in *RdRp*^tg/-^ mice and *RdRp^tg/-^Adar^+/-^*mice. Notably, in *Adar^+/-^* mice alone, we observed decreased ADAR1 protein expression (**Fig. 7F**), as would be predicted from loss of an allele.

Endogenous retroelements are enriched in the 3’ UTRs of ISG mRNAs (68). Despite the increased ISG mRNA expression in *RdRp^tg/-^ Adar^+/-^* animals compared to WT animals (**Fig. 5C**), the proportion of uniquely edited sites that were within an ISG RNA versus those of other genes was not significantly elevated (**Fig. S10**). This was the case despite the double heterozygotes having an increase in total edited genes and sites (**Fig. 7C**). Compared to the WT and double heterozygotes, *RdRp^tg/-^* and particularly *Adar^+/-^* animals had small, though significant, reductions in the proportion of uniquely edited sites falling within an ISG. These results suggest that while ISGs themselves were also edited, the amount of editing within the ISG mRNAs does not account for the observed dysregulation of Adar editing.

In summary, single *Adar* allelic loss coupled with transgenic expression of the picornavirus RdRp synergize to cause an extreme ISG response which produces an interferonopathy remarkably similar to Singleton-Merten syndrome. This work presents a novel model for this disease, which can be further studied to unlock molecular mechanisms driving the pathology. The ADAR-intact model is exceptional for its paradoxical combination of tolerated MDA5 hyperactivity with autoimmunity resistance.

## DISCUSSION

Precise regulation of intracellular RNA duplex sensing is critical for protecting mammals against the Scylla of viral disease while also evading the Charybdis of autoimmunity. Confining sensing to only exogenous dsRNAs via PRRs such as MDA5 is therefore a bedrock feature of healthy antiviral defense. However, since MDA5 is relatively non-discriminating in its ligand preferences (mainly binding to internal segments of longer dsRNA duplexes on the order of hundreds to thousands of nucleotides in length), it has become clear in the past few years that an additional layer of control, ADAR1 p150 editing of host cellular RNAs, is at play. Nevertheless, the biological significance of the millions of A-to-I edits that modify RNA transcripts are incompletely understood.

In the same year in which he published his Nobel-awarded paper on the YF17D yellow fever vaccine that is still used today (69), virologist Max Theiler reported his isolation of Theiler’s virus and noted its similarities to poliovirus (70). Here we identify new features of a mouse model in which we express the dsRNA-synthesizing enzyme of this picornavirus outside the viral context in the tissues of mice, providing a continuous dsRNA stimulus to the MDA5-MAVS pathway. In its strict MDA5-MAVS dependence, the model parallels the exclusively MDA5-dependent pattern for sensing of replicating picornaviruses (1, 5–8). Having previously demonstrated the lack of significant autoinflammatory ‘cost’ to the animals, we found here that RdRp^tg^ mice are further able to resist an autoimmune provocation of induced SLE in the BM12 model (**Fig. 1**).

However, the loss of a single *Adar* allele, which by itself is not discernibly harmful in our and others’ hands (18), breaks the autoinflammation-protected state, resulting in a severe disease with progeric features, characterized by shortened lifespan, stunted growth, premature fur graying, poorly developed teeth, and skeletal abnormalities, extreme ISG elevations, and dysregulated and increased A-to-I editing. In addition to the accentuation of ISG hyper-expression compared to RdRp^tg^ mice, which may be causative in the pathology that emerged, the double heterozygotes hyper-expressed mRNAs for several CC and CXC cytokines (**Fig. S8**), a result that will be interesting to explore further in future studies of these mice. In contrast, type I IFNs and the classical proinflammatory cytokines IL6 and TNFα were not induced to higher levels by addition of the single *Adar* allele knockout.

Among ADAR1 mouse models that produce autoinflammatory pathology, this is the only one with a simple haploinsufficient genotype and a wild type remaining *Adar* allele. All others are either *Adar* ^-/-^ (which is embryonic lethal without an *Mda5* KO rescue) or utilize protein mutants such as *Adar*^P95A^ or *Adar*^E861A^ [see ref. (71) for a comprehensive tabulation of existing mouse models]. Relatedly, we further emphasize that the present model is nucleic acid-driven rather than MDA5 GOF mutation-triggered. As such it is a mouse model in which ‘endogenously’ synthesized RNAs generate an autoinflammatory disease via wild type MDA5, with the source of the dsRNA being a genomically-integrated viral polymerase. Since loss of one *Adar* allele is enough to tilt the immune system in RdRp^tg^ mice to a highly pathogenic autoinflammatory outcome, it can be speculated that in some natural circumstances viral infections and the inflammatory signaling cascades they induce have roles in triggering human autoinflammatory diseases, particularly if they also disturb the finely regulated, complex equilibrium of A-to-I editing. The RdRp^tg/-^ *Adar^+/-^* model complements ADAR1 mutant mouse AGS models, which differ in utilizing mutant ADAR1 proteins (72–74), and which do not incorporate a viral polymerase. The pathological outcome in *RdRp^tg/-^ Adar^+/-^* mice is variably penetrant, with about half of mice severely affected, a feature that is common in human SMS kindreds (55, 62) as well ADAR1 mutant protein mouse models of AGS (24, 73).

In some respects, the pathologies observed in *RdRp^tg/-^ Adar^+/-^* mice resemble the human interferonopathy SMS, with notable exceptions, e.g., the lack of aortic and valvular calcification typically seen in the human syndrome (54, 55). In previous studies phenotypes resembling AGS/SMS were produced by engineering mice to express the MDA5 constitutive activation (GOF) mutants mG821S (75) or huR822Q (76). In mG821S mice, SMS-like features were observed (75) but the model differs in that while it recapitulates skeletal defects, also seen in *RdRp^tg/-^ Adar^+/-^* mice and human SMS cases (56, 57, 59), it did not have the dental deformities or the fur pigmentation abnormality we observed, nor did we see the deteriorating kidney function found in mG821S mice. In the picornavirus RdRp^tg^ model, constitutive activation of WT MDA5 expressed from the endogenous locus by upstream provision of sustained dsRNA production is insufficient and addition of *Adar* haploinsufficiency is needed. It may thus have value as a model of nucleic acid-induced autoimmunity that reflects the overwhelming majority of patients, who have normal RLR proteins. In this regard, the G821S and R822W GOF MDA5 mutant proteins do not bind dsRNA and are always “on”. This may be central to the difference. Subsequent investigation of the pathology observed in *RdR^tg^ Adar* haploinsufficient mice will ideally test a potential role for ZBP1, which has been identified as a main downstream effector of autoinflammation in the context of mutant ADAR1 proteins. ADAR1 acts as an upstream negative regulator of its activation, which can trigger inflammatory signaling and several varieties of regulated cell death (63, 68, 77–80)

Why ADAR1, which mediates the most abundant form of RNA editing in metazoan organisms (81), does not edit RdRp-synthesized RNAs sufficiently to prevent activation of WT MDA5 in the RdRp^tg^ but *Adar^+/+^* mouse is unclear and is an intriguing aspect to investigate in the future. The data indicate that mammals have a means to prevent viral RdRp-synthesized dsRNA from being protected (from sensing) by A-to-I editing, even when the RdRp is expressed completely outside the context of actual viral replication, i.e., without the elaborate replication factory biogenesis central to the life cycles of all positive strand RNA viruses (and also without Vpg protein capping in the case of picornaviruses). How this occurs is a key topic for investigation in the field.

In prior reports of *Ifih1* gene duplication or mice carrying MDA5 GOF mutants, which also have increased expression of ISGs and relative protection against viral diseases, the mice have been shown to be more prone to either spontaneous or triggered autoimmunity (36, 42, 43, 75, 76). Thus, RdRp^tg^ mice may have distinctive tolerance mechanisms as evidenced by their resistance to induced SLE. A component of these tolerance mechanisms may be the increase in overall and effector regulatory T cells (**Fig. 1**), which are key autoimmune suppressor cells (49).

We conclude from our data that correctly regulated ADAR1 editing is a key suppressor of interferonopathic outcomes in these animals. The results also suggest that modulation of ADAR1 activity might potentially be of benefit in some human autoinflammatory diseases. Despite loss of one allele, ADAR1 levels, specifically the levels of ADAR p150 protein were increased equivalently in both *RdRp^tg/-^* and *RdRp^tg/-^ Adar+/-* animals, indicating that disease is not driven by a gene dosage effect in which low ADAR1 levels cause hypo-editing of dsRNA. Consistent with this result, a dysregulation picture emerged in which the overall number of edited sites and edited genes are increased in the double heterozygotes as compared to all other genotypes and are abnormally distributed (**Fig. 7**). The dysregulation could contribute to the severe phenotype observed, but again, the comparison of the degree of p150 elevation with that of *RdRp^tg/-^*mice compels an interpretation that the level of induced ADAR1 protein is not the main driver. For example, abnormally high levels of RNA editing have been reported in SLE patients (82). When we measured the levels of proteins encoded by several uniquely edited ISG transcripts in *RdRp^tg/-^Adar^+/-^*mice, we did not identify an effect of the additional edits. We considered whether increased ISG expression might lead to increased endogenous retroelement dsRNA expression, a possibility suggested by the observed increase in editing of SINES and LINES; in this regard, Zhang et al. showed that such endogenous retroelements are in fact quite enriched in the 3’ UTRs of ISG mRNAs (68). Here we found that multiple ISG mRNAs were indeed A-to-I edited but the proportion of uniquely edited sites that were within an ISG RNA versus those of other genes was not significantly elevated.

ADAR1 activity and editing changes in autoinflammatory diseases that are not directly linked to ADAR1 mutations are still poorly understood, but our data indicate that simple expression of the enzyme at physiological or higher levels does not equate with proper function. It will be worthwhile to study changes in ADAR1 expression or dysregulation of A-to-I editing in interferonopathies and other autoimmune disease not directly linked to ADAR1.

## METHODS

### Mice and nomenclature

RdRp transgenic mice on the C57/BL6J background have been previously described (37). We here use *RdRp^tg/-^*to designate a mouse with one RdRp transgene allele and one normal locus lacking the chromosome 6 transgene insertion, *RdRp^tg/tg^*for transgene homozygotes, and RdRp^tg^, without italics, for the general model. *Adar^+/-^* mice on the C57/BL6 background generated by Hartner et al (20). were obtained from the Jackson Laboratory (MGI 3029789). There is a germline deletion of the gene region spanning exons 7-9, which results in non-functional p110 and p150 proteins (20). Mice were backcrossed for two generations onto our colony before use and genotypes were confirmed using protocols suggested by the Jackson Laboratory. CD45.1^+^ congenic marker line mice were obtained from the Jackson Laboratory (mouse strain: 002014) and crossed onto the BM12 mouse line (Jackson Laboratory, 001162). Presence of the CD45.1 gene was confirmed via TransnetYX genotyping service (Ptprc-2 Mut probe) and confirmed with flow cytometric staining of splenocytes. BM12 genotyping was confirmed as described (45) in *Ifih1* knockout mice on a B6.J background were obtained from Jackson labs (strain 015812). As much as possible, littermate controls were used for direct comparison between groups. All experiments use a mix of male and female mice for analysis. All experiments were performed in accordance with the IACUC guidelines and approved procedures for the University of Colorado, Anschutz Medical Center (IACUC protocol no. 00116). BM12 initiation of lupus like disease was done as described (45). Briefly, 1×10^8^ splenocytes isolated from CD45.1^+^ BM12 in mice were injected intraperitoneally into WT(CD45.2) or *RdRp^tg/tg^* (CD45.2) mice. Single cell solution in PBS or PSB alone (sham) was used for injection in 250 μL total volume. Mice were euthanized at 14 days post injection for disease evaluation.

### Animal Imaging, µCT analysis, Intraocular pressure determinations

Femurs were evaluated for length, cortical bone structure and trabecular microarchitecture using a Zeiss Xradia Versa X-ray microscope-520 (μCT, Zeiss, Dublin, CA, USA; 80 kVp, 4x objective, isotropic voxel size of 4 μm). Regions of interest were located at the femur mid-diaphysis (1 mm tall) and the distal femur metaphysis (starting at 600 μm proximal to the epiphyseal line and extending 1,000 μm proximally). Trabecular and cortical bone were segmented and analyzed using Dragonfly Pro and Bone Analysis software (Object Research Systems, Montreal, Canada) according to established guidelines (83). Trabecular bone parameters included bone volume fraction (Tb.BV/TV), trabecular number (Tb.N), trabecular thickness (Tb.Th), and trabecular separation (Tb.Sp). Cortical bone measures included cortical bone volume fraction (Ct.BV/TV), total bone mineral density (TMD), and cortical porosity (Ct.Po). The bending moments of inertia (Imax and Imin) which contribute to structural stiffness under bending were calculated. Hydrated femurs were evaluated using three-point bending to failure as described (84, 85) to determine mechanical properties (e.g., stiffness and maximum load) using a MTS Insight II benchtop tester (MTS Corp., Minneapolis, MN; 250 N load cell, 5 mm/min deflection rate, 7 mm span width) and analyzed using custom MATLAB code, Mathworks, Natick, MA, USA). Material properties (e.g., modulus, ultimate stress) were calculated using standard equations derived from engineering beam theory (84). Intraocular pressures were measured with a handheld ocular pneumotonometer (Model 30 Classic, Medtronic) as described (86).

**See Supplemental Methods for**

a. Cell culture and siRNA delivery.
b. Bioinformatics and Computational Methods.
c. Flow Cytometry
d. Quantitative PCR and Immunoblotting
e. ELISAs.
f. Calcium Quantification
g. Histopathology methods and scoring

## ACKNOWLEDGMENTS

Funding was provided by National Institutes of Health Avant-Garde grant DP1DA043915 (E.M.P.), a John H. Tietze Foundation grant (E.M.P.), and an NSF grant 1707065 (V.L.F.). We thank C. Radomile and J. Chen for technical assistance with animal experiments, the University of Colorado Denver School of Medicine (UCDSOM) Division of Allergy and Clinical Immunology Flow Cytometry Core for use of the LSRII instrument, the UCDSOM Research Histology Core Laboratory for tissue processing (RRID: SCR_021994), and the CU at Boulder Materials Instrumentation and Multimodal Imaging Core Facility (RRID: SCR_019307) for X-ray microscopy and micro-computed tomography imaging.

## Supplemental Figure Legends

**Fig. S1.**
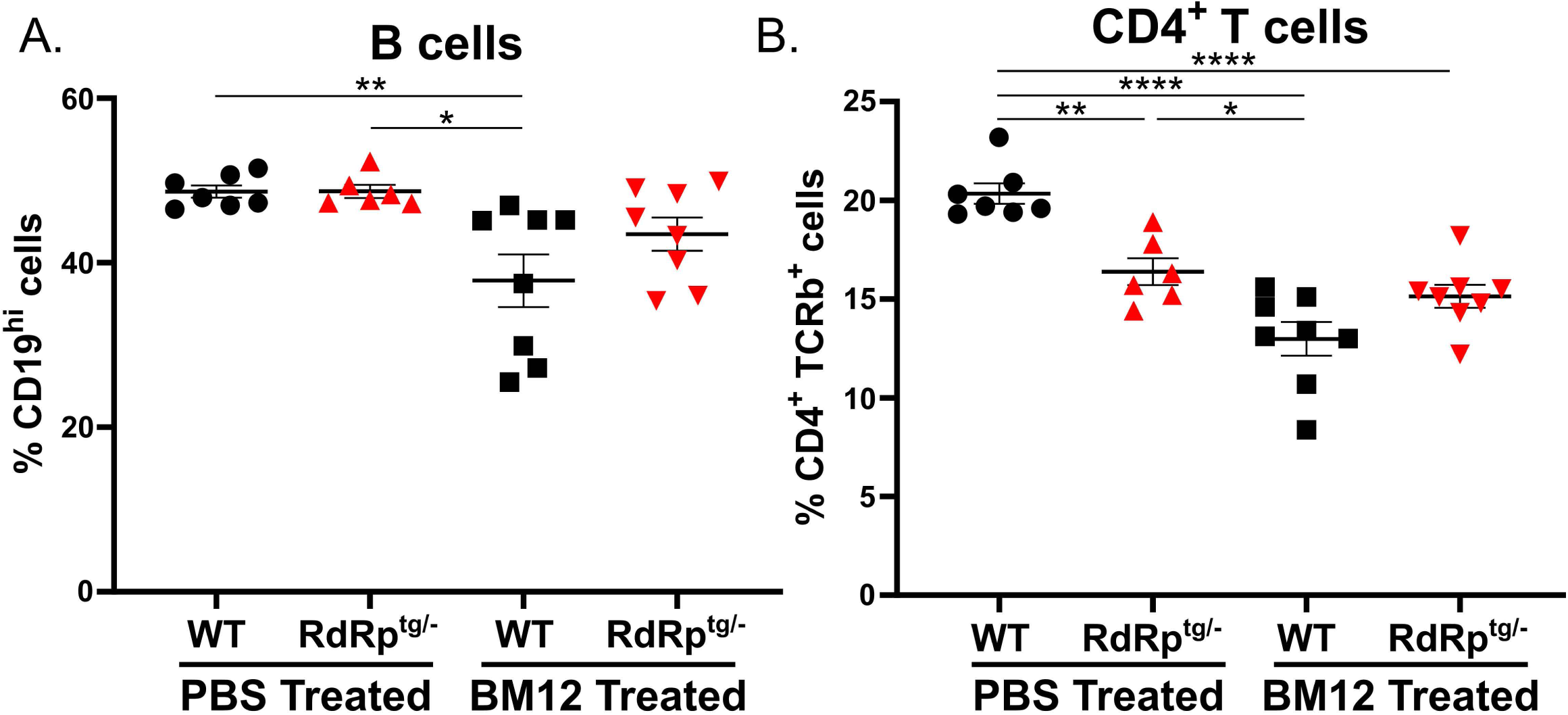
Overall B cell and T cell populations are marginally changed during BM12 transfer. Single cell, RBC-lysed splenocyte suspensions were made from sham or BM12 transferred animals for flow cytometry analysis. **(A)** Overall B cell populations in animals as defined by cd19 expression. **(B)** Overall CD4^+^ T cell population in the spleens of experimental animals, as defined by TCRβ and CD4 expression. Data were analyzed using a one-way ANOVA followed by a Tukey tests where * = p < 0.05, ** = p < 0.01, *** = p < 0.001, **** = p < 0.0001. n = 7, 6, 8, 8 for WT (PBS), *RdRp^tg/-^* (PBS), WT (BM12), and *RdRp^tg/-^* (BM12), respectively. Data represent individual animals with the mean and s.d. shown as bars.

**Fig. S2.**
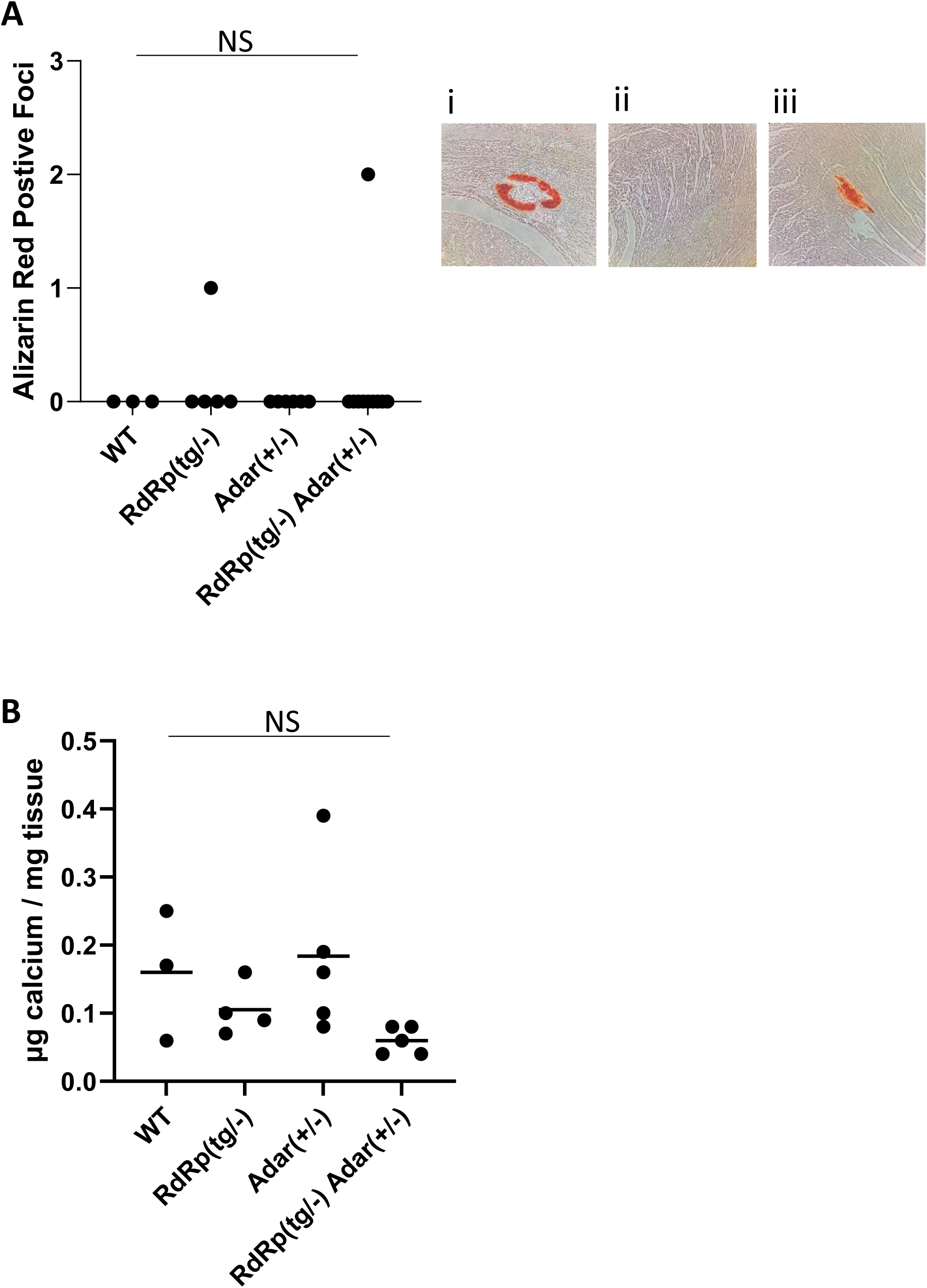
Analysis of organ calcification. Flash frozen brain and formalin-fixed paraffin embedded heart were collected from 6-week-old mice. **(A)** Heart samples were stained with Alizarin Red and calcium depositions were manually counted and shown as Alizarin Red foci per animal (n = 3, 5, 5, 10 for WT, *Adar^+/-^*, *RdRp^tg/^*, *RdRp^tg/-^ Adar^+/-^*, respectively). Representative heart section images are shown to the right: (i) positive control heart provided by UCDSOM Research Histology Core Laboratory; (ii) RdRp^tg/-^ Adar^+/-^ heart without Alizarin red foci; (iii) RdRp^tg/-^ Adar^+/-^ heart with a positive focus. **(B)** Calcium was quantified from brain tissues (n = 3, 4, 5, 5). Data were analyzed using a one-way ANOVA followed by a Tukey tests; there were no significant differences between the groups. ns: not significant.

**Fig. S3.**
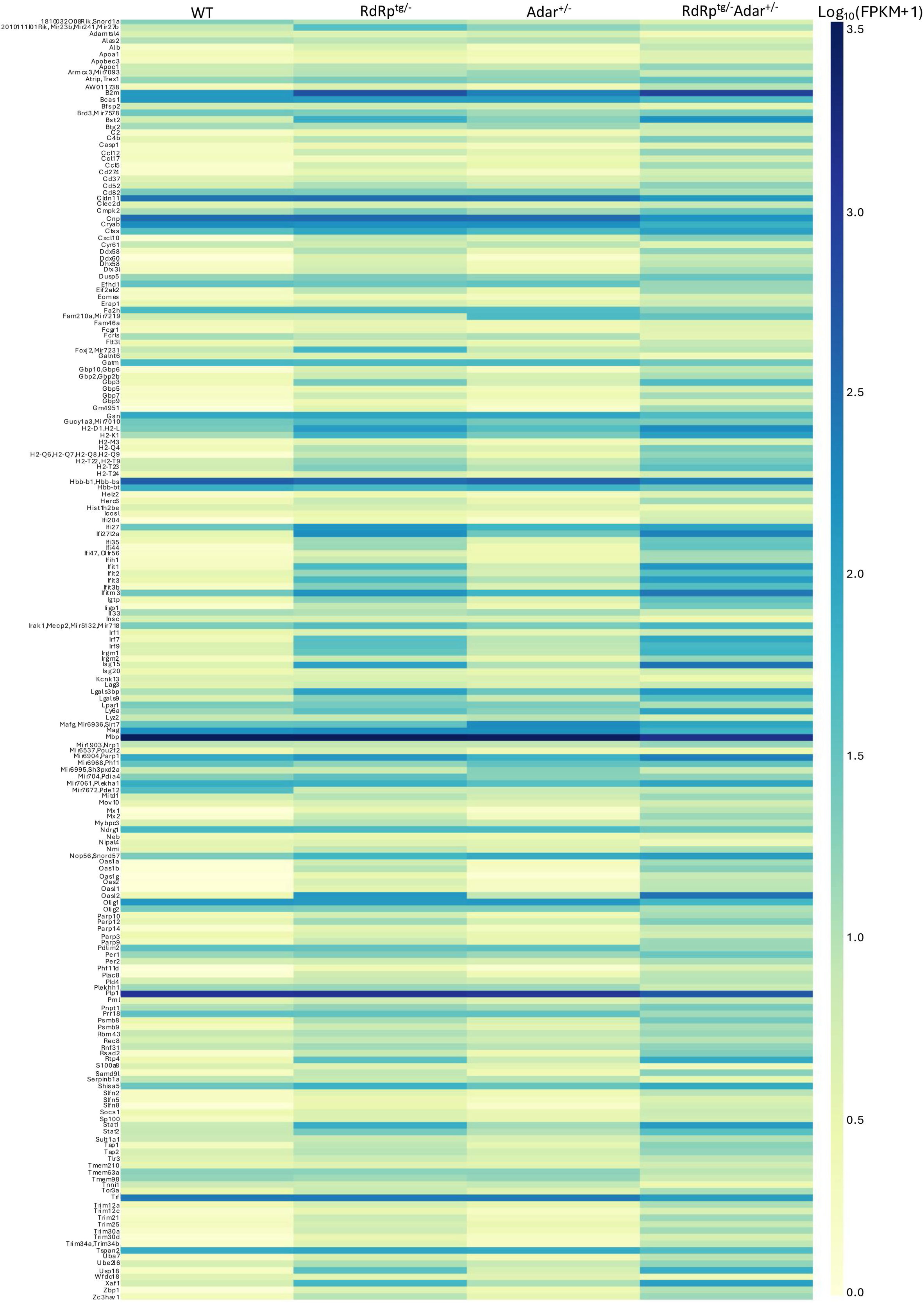
Heatmap of differentially expressed ISGs across the four genotypes. Enlarged image of Fig. 5B with gene names displayed. FPKM values were log transformed with 1 pseudocount to facilitate visualization.

**Fig. S4.**
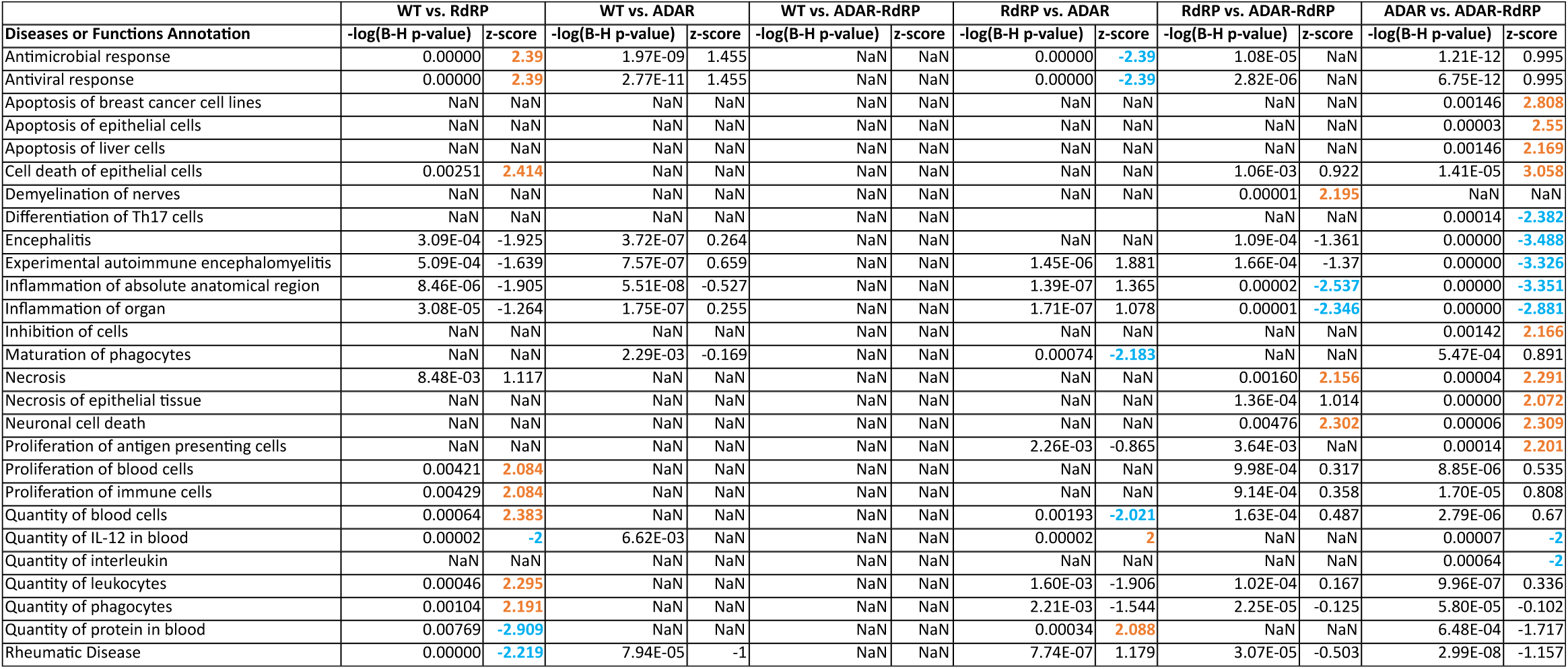
Predicted upstream regulators. Predicted upstream signaling regulators as determined by IPA in various genotype comparisons between WT, *Adar^+/-^*, *RdRp^tg/-^,* and *Adar^+/-^ RdRp^tg/-^* mice where red/green indicates whether the gene itself is upregulated/downregulated in the RNA-seq and orange/blue indicates predicated activation/inhibition of the molecular pathway it regulates.

**Fig. S5.**
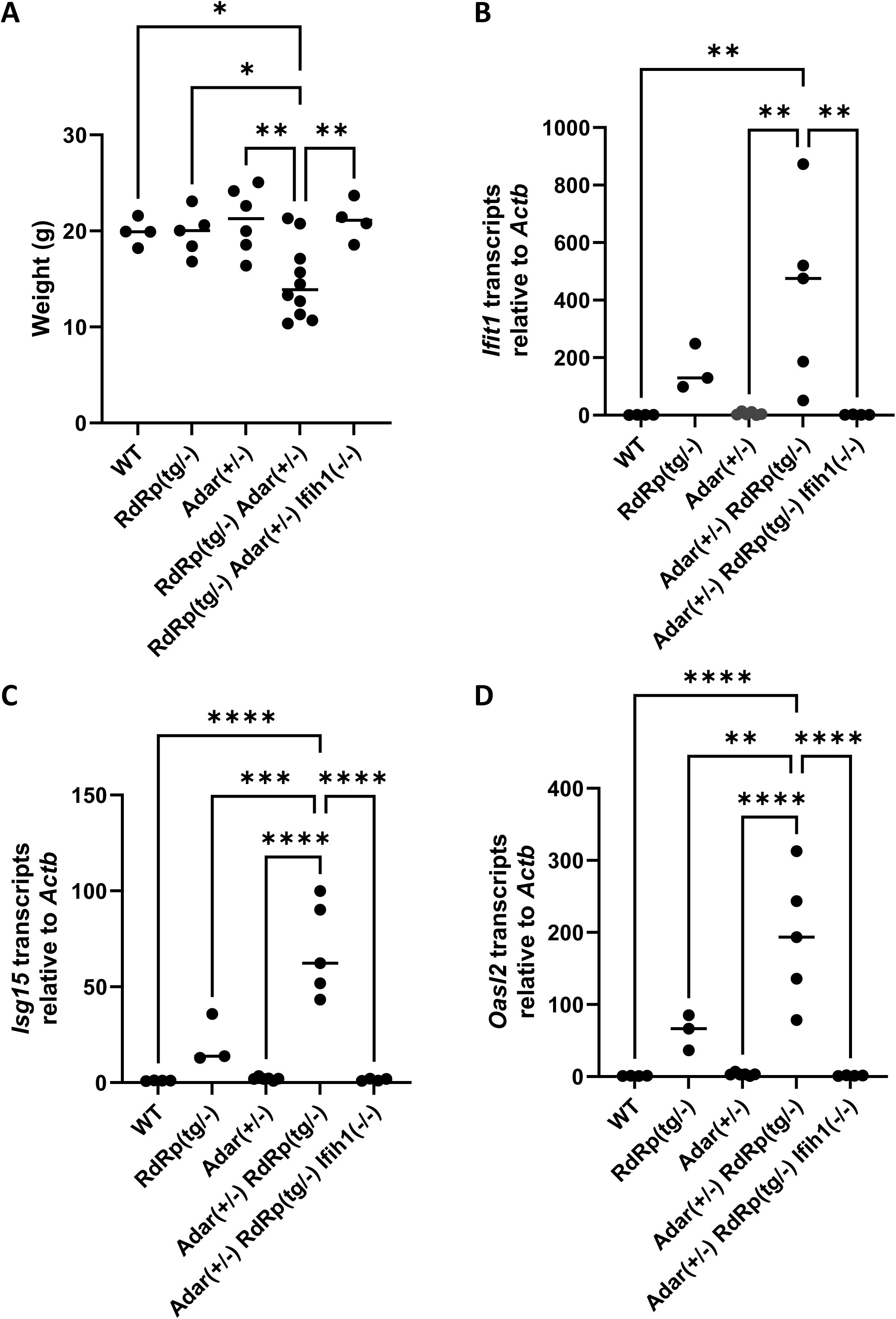
Loss of MDA5 rescues runting phenotype and abolished ISG upregulation. 6- week-old animals were weighed **(A)** and RNA was isolated from whole-brains for qPCR analysis of the ISGs Ifit1 **(B)**, Isg15 **(C)**, and Oasl2 **(D)**. Data were analyzed using a one-way ANOVA followed by a Tukey tests where * = p < 0.05, ** = p < 0.01, *** = p < 0.001, **** = p < 0.0001. For (A), n = 4, 5, 6, 10, and 4 for WT, *Adar^+/-^*, *RdRp^tg/^*, *RdRp^tg/-^ Adar^+/-^*, and *RdRp^tg/-^ Adar^+/-^ Ifih1^-/-^*, respectively. For (B-D), n = 4, 3, 5, 5, and 4. Data points represent individual animals with the mean for each group shown as a bar.

**Fig. S6.**
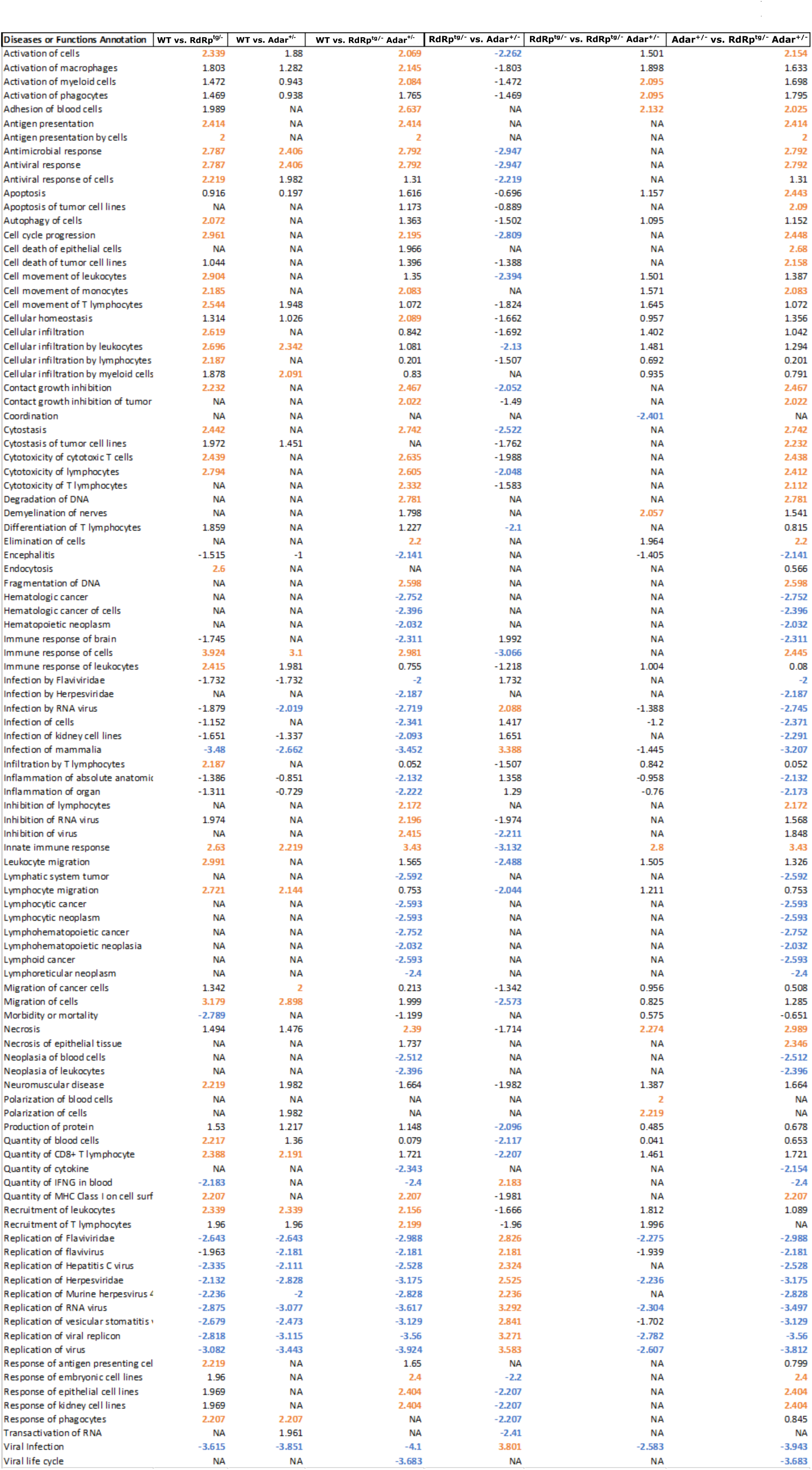
Disease and Biological Functions. Enriched diseases and biological functions across comparisons of gene expression changed between WT, *Adar^+/-^*, *RdRp^tg/-^,* and *Adar^+/-^ RdRp^tg/-^* mice. Orange indicated that the disease or biological function is predicted to be enriched for and blue indicated is predicted to be inhibited.

**Fig. S7.**
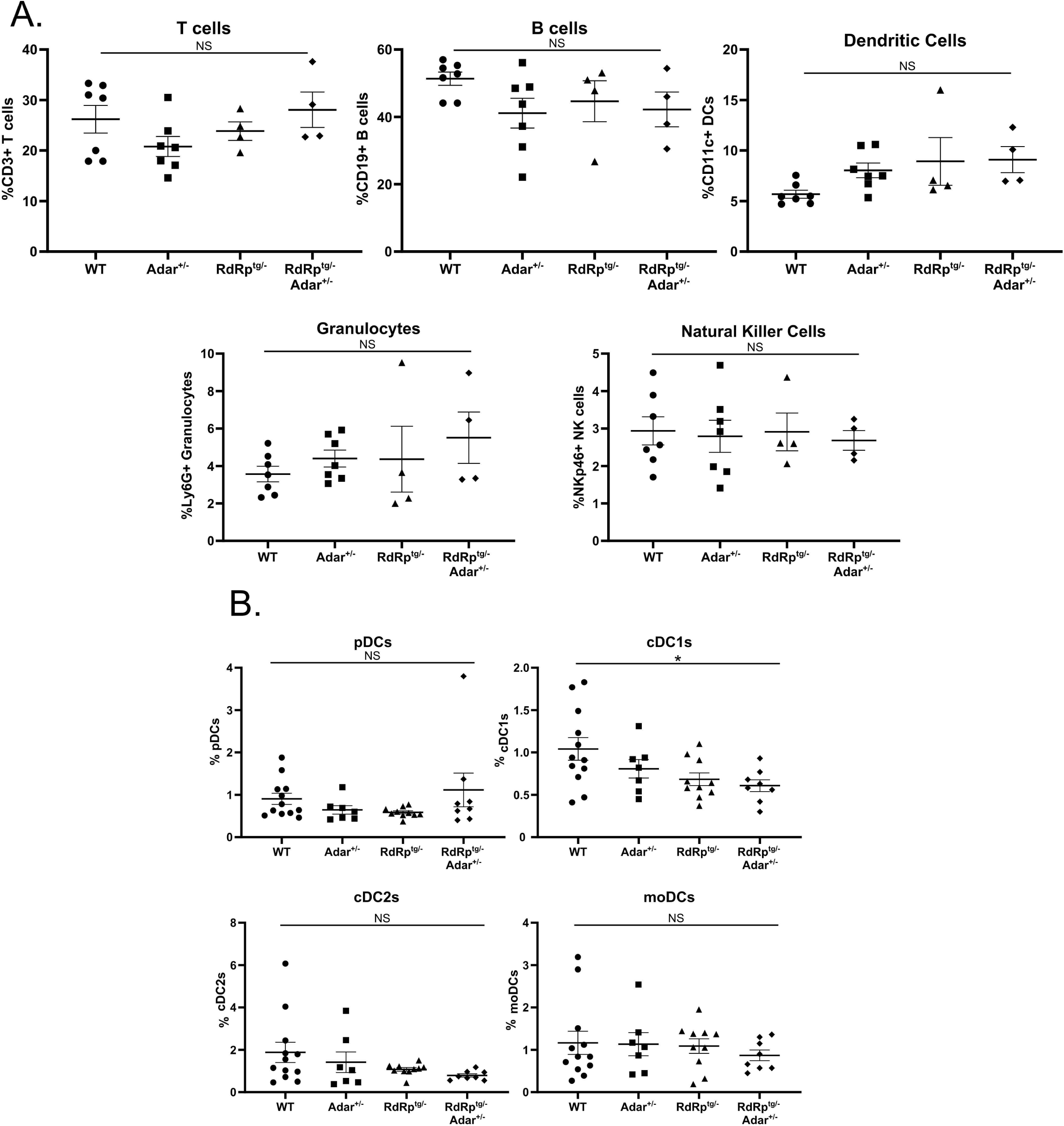
Certain immune cell populations are unchanged in *RdRp^tg/-^ Adar^+/-^* mice. Splenocytes from 4-5 week old mice were harvested and single cell, RBC-lsysed solutions were made for FACS analysis. **(A)** T cells, B cells, DCs, granulocytes, and NK cells show no difference across groups measured. n = 7, 7, 4, and 4 for WT, *Adar^+/-^*, *RdRp^tg/^*, and *RdRp^tg/-^ Adar^+/-^*, respectively. **(B)** DC subsets including pDCs, cDC1s, cDC2s, and moDCs show minor or no difference in populations across groups. n = 12, 7, 10, and 8 for WT, *Adar^+/-^*, *RdRp^tg/^*, and *RdRp^tg/-^ Adar^+/-^*, respectively. ll data was analyzed using a one-way ANOVA followed by a Tukey test to determine significance. * = p < 0.05. Data represent individual animals, graphs show means and s.d.

**Fig. S8.**
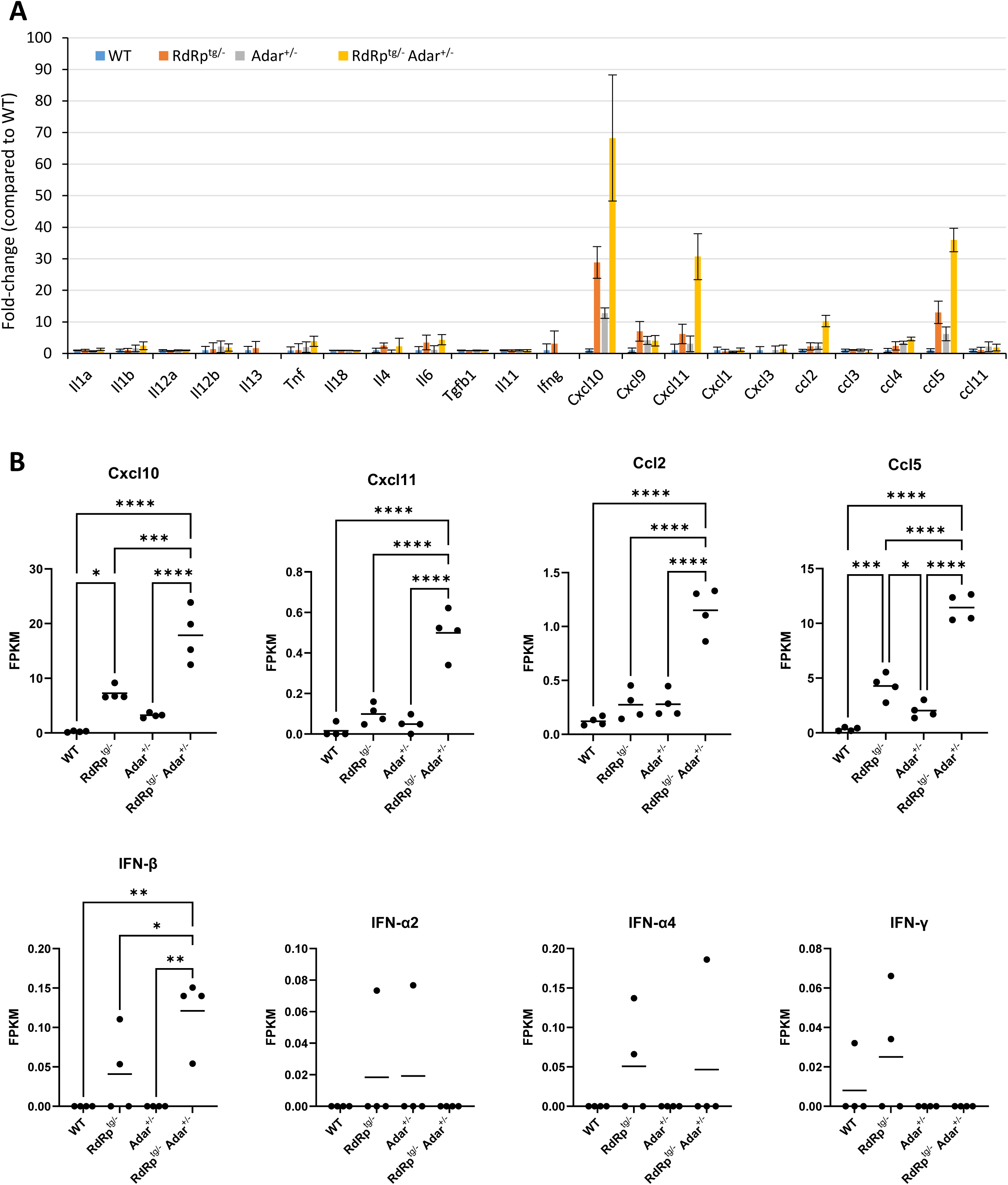
Chemokine, cytokine and IFN expression changes. **(A)** RNA-seq-derived gene expression changes were determined for chemokines and cytokines and are shown as the average fold-change compared to wild-type mice, +/- s.d. **(B)** Individual animal FPKM values for cxcl10, cxcl11, ccl2, ccl5, IFN-β, IFN-α2, IFN-α4, and IFN-γ were graphed separately. Data were analyzed using a one-way ANOVA followed by a Tukey test to determine significance. No significant changes were observed for IFN-α2, IFN-α4, and IFN-γ. * = p < 0.05, ** = p < 0.01, *** = p < 0.001, **** = p < 0.0001.

**Fig. S9.**
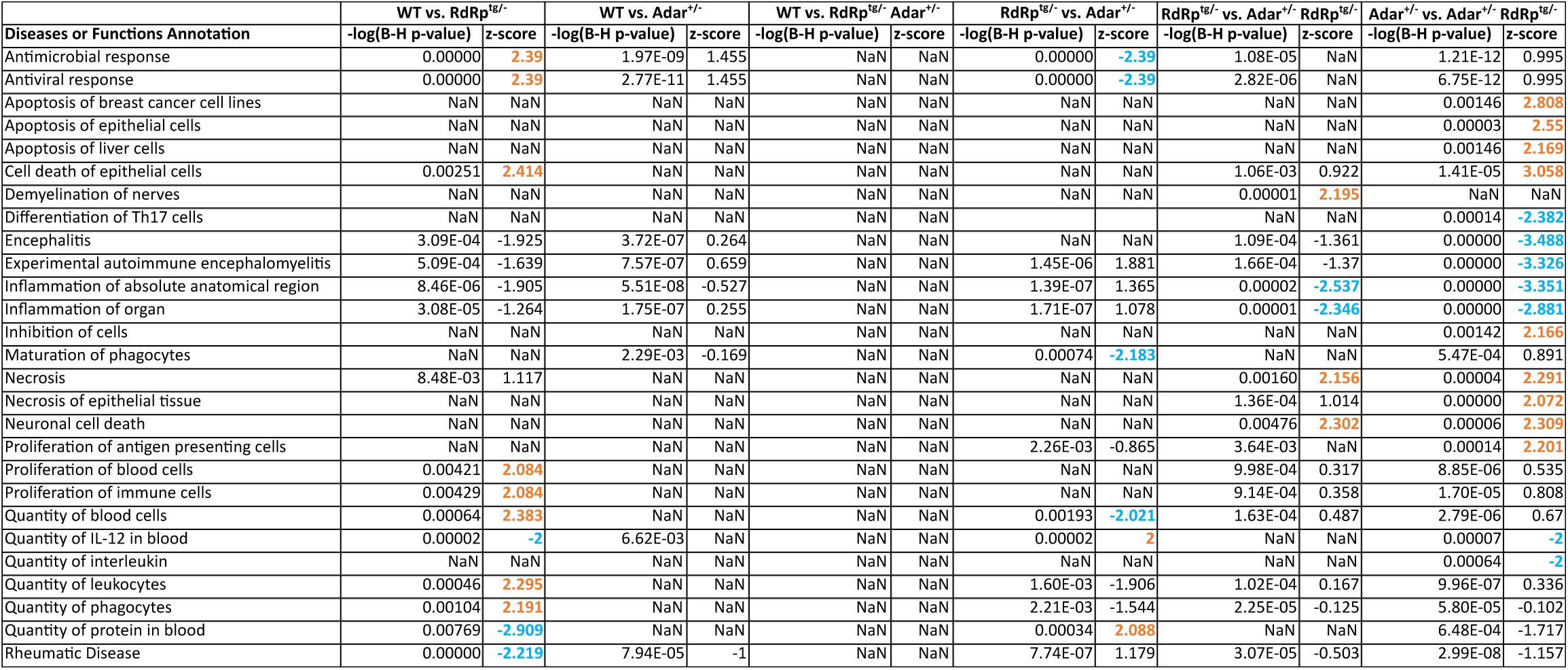
Diseases and Biological Functions enriched for in A-to-I edited genes. Genes with A-to-I edit sites and RNA-seq-derived expression changes were analyzed using IPA to determine enriched diseases and biological functions across genotype comparisons (WT, *Adar^+/-^*, *RdRp^tg/-^,* and *Adar^+/-^ RdRp^tg/-^*). Orange indicates that the disease or biological function is predicted to be enriched for or activated and blue indicates is predicted to be inhibited.

**Fig. S10.**
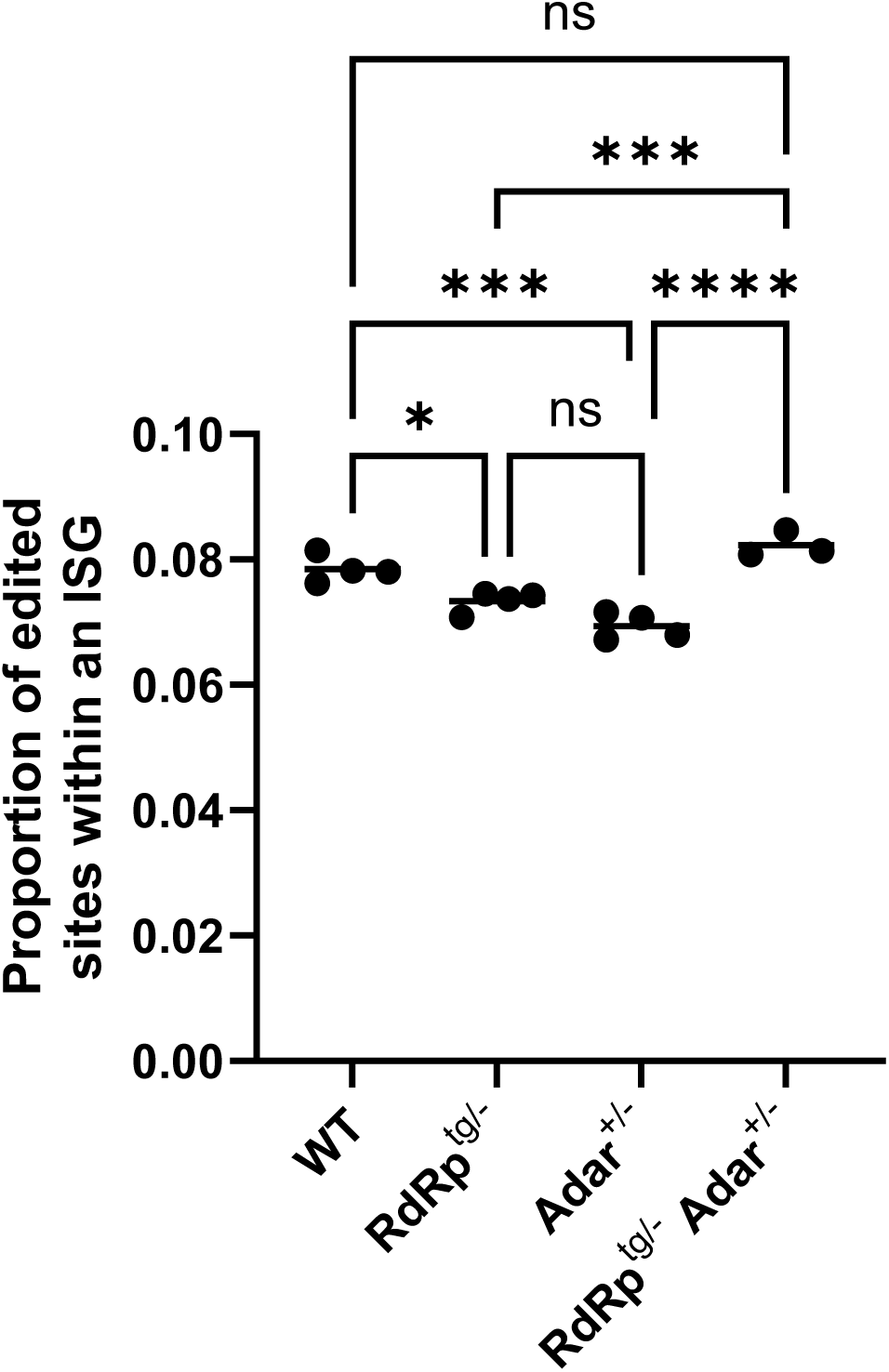
Proportion of edited sites in ISG RNAs as a fraction of all uniquely edited sites per animal. Sites of A-to-I editing that were within an ISG were identified and the proportion of ISG edited sites out of all unique edited sites per animal was calculated. Bars indicate group means, dots represent individual animals, and the error bars are s.d. Data were analyzed using a one-way ANOVA followed by a Tukey tests where * = p < 0.05, *** = p < 0.001, **** = p < 0.0001, ns = not significant.

## Supplemental Methods, Miller et al

### PCR primers and antibodies

#### a.#Primers

##### Mus musculus

Gapdh: 5’-CATGGCCTTCCGTGTTCCTA-3’ & 5’-CTATGTTTCCCGCGGCACGTCAGATCCA-3’

Ifit1: 5’-TTTGTTGTTGTTGTTGTTCGTT-3’ & 5’-GCAGGAATCAGTTGTGATCT-3’

ISG15: 5’-TGGTACAGAACTGCAGCGAG -3’ & 5’-CAGCCAGAACTGGTCTTCGT -3’

Oasl2: 5’-CAAACAAACAAACAAACCCTCTC-3’ & 5’-TCAAGGTGTCACTCTGCAT-3’

Adar p150 5’-GGCCTTGCCGGCACTATGTCTC-3’ & 5’

GCGGGTATCTCCACTTGCTATGCTC-3’

Adar p110: 5’-GGTGGAAGACTACGCGTTGGGAC-3’ & 5’ ACGACTGTGTCTGGTGAGGGAACAC-3’

IFNβ: 5’-CACAGCCCTCTCCATCAACTATAAGCAG-3’& 5’-

CTGTAGGTGAGGTTGATCTTTCCATTCAG-3’

##### Homo sapiens

GAPDH: 5’-TGGACCTGACCTGCCGT-3’ & 5’-TGGAGGAGTGGGTGTCGC-3’

ISG15: 5’-GTGGACAAATGCGACGAACC-3’ & 5’-ATTTCCGGCCCTTGATCCTG-3’

OASL: 5’-GGATCTTCTCCCACACTCACATCT-3’ & 5’-CACCATCAGGATTCTTCACGAA-3’

#### b.#Antibodies

Rabbit polyclonal anti-ADAR1 (1:1,000 dilution; Cell Signaling, 14175)

Mouse anti-tubulin (1:5,000 dilution; Sigma, T5168)

Rabbit anti-Rig-I: mouse tissue blots (1:1,000 dilution, Cell Signaling, 3743)

Mouse anti-ADAR1 (1:500 dilution, Santa Cruz Bio, sc-73408)

Rabbit anti-actin (1:5,000 dilution, Cell Signaling, 4967)

Goat anti-rabbit secondary antibody, horseradish peroxidase -conjugated (1:5,000 dilution; Calbiochem, 401393)

Goat anti-mouse secondary antibody horseradish peroxidase-conjugated (1:5,000 dilution; Invitrogen, 31430)

Rabbit anti-ZBP1/DAI (1:1,000 dilution; Novus Biologicals NBP1-76854)

Rabbit anti-ISG15 (1:1000, Cell Signaling Technologies Cat # 2743)

### Quantitative PCR

For qPCR analysis of mouse samples, RNA was extracted from mouse tissue using Trizol reagent (Ambion) according to manufacturer instructions. Reverse transcription PCR (RT-PCR) for cDNA synthesis was conducted using Maxima H Minus First Strand cDNA Synthesis Kit (Thermo Scientific) following manufacturer instructions. QPCR analysis for quantification of cellular transcripts was done using 2X qPCR Green Master Mix (APEX). For qPCR analysis from human cell lines, RNA extractions were done using an RNAeasy kit (Qiagen) according to kit protocol. RT- and qPCR were performed as described above. Transcripts were quantified using the ΔΔC_T_ method relative to *Gapdh* or β-Actin (*Actb*) transcripts.

### Immunoblotting

Cells were homogenized in RIPA buffer and protein content was determined by Bradford reaction (Bio-Rad). 30 μg total protein was loaded per well prepared in 6X reducing SDS sample buffer (Bio-Rad, boil for 5-10 minutes prior to loading). Samples were run on 10% agarose gels (Bio-Rad) and transferred to polyvinylidene difluoride membrane (Bio-Rad).

Samples were blocked in 5% milk in tris-buffered saline containing 0.1% Tween20 (TBS-T). Immobilon Crescendo Western HRP substrate (Millipore) was used for visualization of bands and imaged on a ChemiDoc XRS+ Imaging System (Bio-Rad).

### Cell culture and siRNA delivery

Triplicate biological replicates were used with three technical replicates each. A549 cell lines (ATTC, CCL-185) were maintained in Dulbecco’s Modified Eagle’s Medium (Corning) containing 10% fetal bovine serum and penicillin/streptomycin/L-glutamine (Corning, 100x solution). Doxycycline-inducible RdRp-expressing cells were described previously (1). Two different siRNAs were used in combination for *ADAR* knockdown and were purchased from ThermoFisher Scientific (Catalog # 4390824, IDs s1008, s1009, previously validated). Negative control (ThermoFisher) or anti-*ADAR* siRNA were transfected into cells (50 pmol per well in a 6-well plate, cells seeded at >50% confluence) using lipofectamine RNAimax reagent (ThermoFisher) according to manufacturer instructions. Cells were washed 6 hours post transfection and dox (2 μg/mL) was added at this time. At 48 hours post dox addition, cells were lysed for protein and RNA analysis.

### Bioinformatics and Computational Methods

We used a combination of RNA sequencing and exome sequencing to evaluate transcriptome-wide patterns of RNA gene expression and ADAR-mediated A-to-I RNA editing. Specifically, we sequenced four biological replicates from each of the four genotypes for RNA-seq (WT-*Adar^+/+^*, WT-*Adar^+/-^*, *RdRp^tg/-^ Adar^+/+^*, and *RdRp^tg/-^Adar^+/-^*) and a single sample from each genotype for exome sequencing. Exome sequencing was used to establish a baseline level of germline genetic variation within our mouse colony and was used as background for identifying RNA-edited sites. Library preparation and sequencing for single-end RNA-sequencing and paired-end exome sequencing were performed by BGI (Beijing Genomics Institute). The quality of raw reads was assessed using FASTQC v.0.11.5. We used Trimmomatic v.0.27 to filter out low quality reads that had a mean Phred quality score less than 20. Read quality was then reassessed with FASTQC. We used Hisat2 v.2.1.0 to map reads to the mm10 genome using default single-end parameters for RNA-seq and included the --no-mixed --no-discordant flags for the paired-end exome sequencing. Next we used Samtools v.1.3.1 to convert the sam output files into bam format. Then we used PicardTools v.1.119 to add read groups, remove PCR duplicates, and coordinate sort the bam files. To evaluate RNA-seq gene expression patterns across the four genotypes, we first used Stringtie v.1.3.6 to quantify transcripts abundance, and Stringmerge to merge the output Stringtie GTF files. CuffDiff2 was to identify significantly differentially expressed genes (DEGs) with an FDR of 0.05 and Benjamini-Hochberg multiple test correction. In order to reduce false positives, we also required fragments per kilobase per million reads mapped (FPKM) to be at least four, and at 2-fold differences in expression because of the massive gene expression differences observed in(1, 2). We used CummeRbund and Plotly to visualize our results. The Interferome Database was used to identify which DEGs were known ISGs. Finally, we used the Canonical Pathways and Diseases and Biological Functions analyses in IPA to determine which molecular pathways and biological functions were enriched among our conditions and to predict whether they were likely to be activated or inhibited based on gene expression patterns in our dataset. To identify RNA-edited sites, we used REDItools to call SNPs that were present in the RNA-seq, but not present in the exome sequencing. Because the A-to-I editing performed by ADAR is incorporated as A-to-G changes by sequencers, we looked for differences in A-to-G changes across conditions. We required a minimum of 10x coverage and that each putative RNA-editing SNP be sequenced at least twice to reduce the risk of including sequencing errors. First, we used Samtools mpileup to call SNPs in the exome sequencing data. We used REDItoolsDeNovo to call SNPs A-to-I edited sites in the RNA-seq requiring at least 10x coverage and that each SNP was sequenced at least twice. Putative germline SNPs identified in the exome sequencing were subtracted from the RNA SNPs. The resulting SNP tables were converted into bed format and sorted by coordinates using BedTools and we filtered SNPs to those present in all four biological replicates. We then annotated the SNPs using Homer annotatePeaks.pl to determine the genomic features in which SNPs occur. A-to-I editing patterns were compared across the four genotypes in two ways and visualized using SUMO, 1) whether editing occurred in the same vs. different genomic features, allowing editing sites to occur at different locations within that feature, and 2) whether editing occurred at identical sites. Finally, we used IPA to identify significantly enriched molecular pathways and biological functions associated with edited genes among the 4 genotypes. The RNA-seq and whole exome sequencing datasets have been deposited to the NCBI sequence read archive (SRA SUB14698103).

### Flow Cytometry

Mice were euthanized by CO_2_ asphyxiation and subsequent cardiac puncture. Spleens were excised and placed in 15 mL conical tubes containing 3 mL of DMEM containing FBS and antibiotics. Spleen was manually dissociated with a pipette tip and pipetting, then passed through a 40 µM nylon cell strainer. 100 µL from each sample were combined and used for the isotype and unstained controls. Cells were gated to exclude debris, doublets, and dead cells based on viability dye staining. Cell population abundance: samples had at least 1×10^6 cells sorted in the live population and cell types are shown as percent of live cells for all flow cytometry data. Each cell type described had a minimum of about 0.5% meaning all cell types described were observed at a frequency of about 5,000 or more per sample. Analysis of cellular populations was conducted as previously described (3). For staining of splenocyte populations, cells were red blood cell depleted as described (3). For analysis including FoxP3 staining, intracellular transcription factor staining was done using Foxp3/Transcription Factor Staining Buffer Set (eBioscience) according to manufacturer instructions for use in a 96-well plate. DC-specific staining panel has been previously described (3). T_reg_ staining panel consists of the following antibodies/cellular stains: fixable viability dye eFlour780 (eBioscience, 65-0865-14), PE anti-mouse FoxP3 (BioLegend, 126404), PerCP-Cy5.5 anti-mouse CD4 (BioLegend, 100434), Brilliant Violet 421 anti-mouse CD25 (BioLegend, 101923), APC anti-mouse CTLA-4 (BioLegend, 106310), Brilliant Violet 711 anti-mouse GITR (BD Bioscience, 563390), Brilliant Violet 605 anti-mouse ICOS (BioLegend, 313538), Brilliant Violet 480 anti-mouse CD44 (BD Bioscience, 566116) and FITC anti-mouse CD62L (BioLegend, 104406). For analysis of B cells (germinal center and plasmablast) the following antibodies and cellular stains were used: fixable viability dye eFlour780 (eBioscience, 65-0865-14), Brilliant Violet 421 anti-mouse CD138 (BioLegend, 142507), PE anti-mouse Fas (BD Bioscience, 554258), Alexa Fluor 488 anti-mouse GL-7 (BioLegend, 144612). For T_FH_ cell analysis and CD45.1 engraftment, the following antibodies and cellular stains were used: fixable viability dye eFlour780 (eBioscience, 65-0865-14), PerCP-Cy5.5 anti-mouse CD4 (BioLegend, 100434), Brilliant Violett 711 anti-mouse TCRβ (BioLegend, 109243), PE anti-mouse PD-1 (BioLegend, 135205), APC anti-mouse CXCR5 (BioLegend, 145506), and Brilliant Violet 421 anti-mouse CD45.1 (BioLegend, 110732). BST-2 was detected with Pacific Blue anti-mouse BST-2 (BioLegend, 127108). Gating for B cell and T_FH_ cell analysis was described in ref (4). Isotype staining for each panel was done in parallel.

Gating was set based on fluorescence minus one controls. Cells were gated to exclude debris, doublets, and dead cells based on viability dye staining. Gating for DC-subtype determination was described in ref. (3). All data was acquired on an LSRII (BD Biosciences) and analysis was done using FlowJo software.

### ELISAs

Mouse serum was used for ANA detection via ELISA. For detection of anti-dsDNA antibodies, calf thymus DNA (Sigma, D4522) was digested with S1 nuclease (ThermoFisher, EN0321) to remove any single stranded DNA (ssDNA). DNA was ethanol precipitated and resuspended in nuclease free water. Freeze-thaw cycles were avoided to prevent creation of new ssDNA. Nunc MaxiSorp ELISA plates (Sigma) were coated with Poly-L Lysine (0.01% final, Millipore Sigma) prior to use. Plates were coated overnight with dsDNA (3 μg/mL) in PBS at room temperature and blocked with BSA in PBS (3% final, Sigma). For a standard, anti-dsDNA (Abcam, ab27156) was used between the range of 1 μg/mL – 1.37 ng/mL (3-fold dilutions) and serum samples were initially diluted 1:10 and then as needed. For anti-smAg ELISAs, plates were coated with Sm Ag (3 μg/mL, Genway Biotech, GWB-CA4A65) in PBS overnight. Blocking was done in 1% BSA/PBS. For standard, anti-SmAg (Novus Biologicals, NB600-546) was used between the range of 1 μg/mL – 15.63 ng/mL (2-fold dilutions) and serum samples were diluted 1:10 before use and then as needed. For both ELISAs, standard and serum samples were diluted in sample buffer (0.5% Tween20, 1% BSA in PBS) and plates were washed 3-5 times between incubations with wash solution (0.5% Tween20 in PBS). Plates were incubated with samples/standard for 2 hours at 37°C. Detection antibody used was goat anti-mouse IgG peroxidase (1:10,000 dilution; Sigma, A2554) and peroxidase was developed used TMB ELISA HRP Substrate kit (Seracare) and Sulfuric acid (4N, Fisher Scientific) according to manufacturer instructions.

### Calcium Quantification

Whole-brain tissue was flash frozen at necropsy, homogenized by sonication on ice in 4x volumes of calcium assay buffer (Calcium Assay Kit, Abcam ab102505), and free Ca^2+^ ions were quantified following the manufacturer’s methods.

### Histopathology

Briefly, approximately 2 cm thick tissue sections of 5 week old mice were fixed in 10% formalin for 24 hours then washed twice and suspended in 70% ethanol. This age was chosen as by this point the mice already showed notable changes in growth but had not yet begun to succumb to disease related mortality. Tissues were paraffin embedded, cut and stained for Alizarin Red or H&E by our Research Histology Core Laboratory. H&E-stained slides were analyzed and scored by the IDEXX BioAnalytics histopathology group and are shown in the figure below. Grading was: 0 no change from WT, 1 minimal, 2 mild, 3 moderate, 4 marked changes. All scores were zero except for one RdRp^tg^ kidney which was scored 1.

**Figure.**
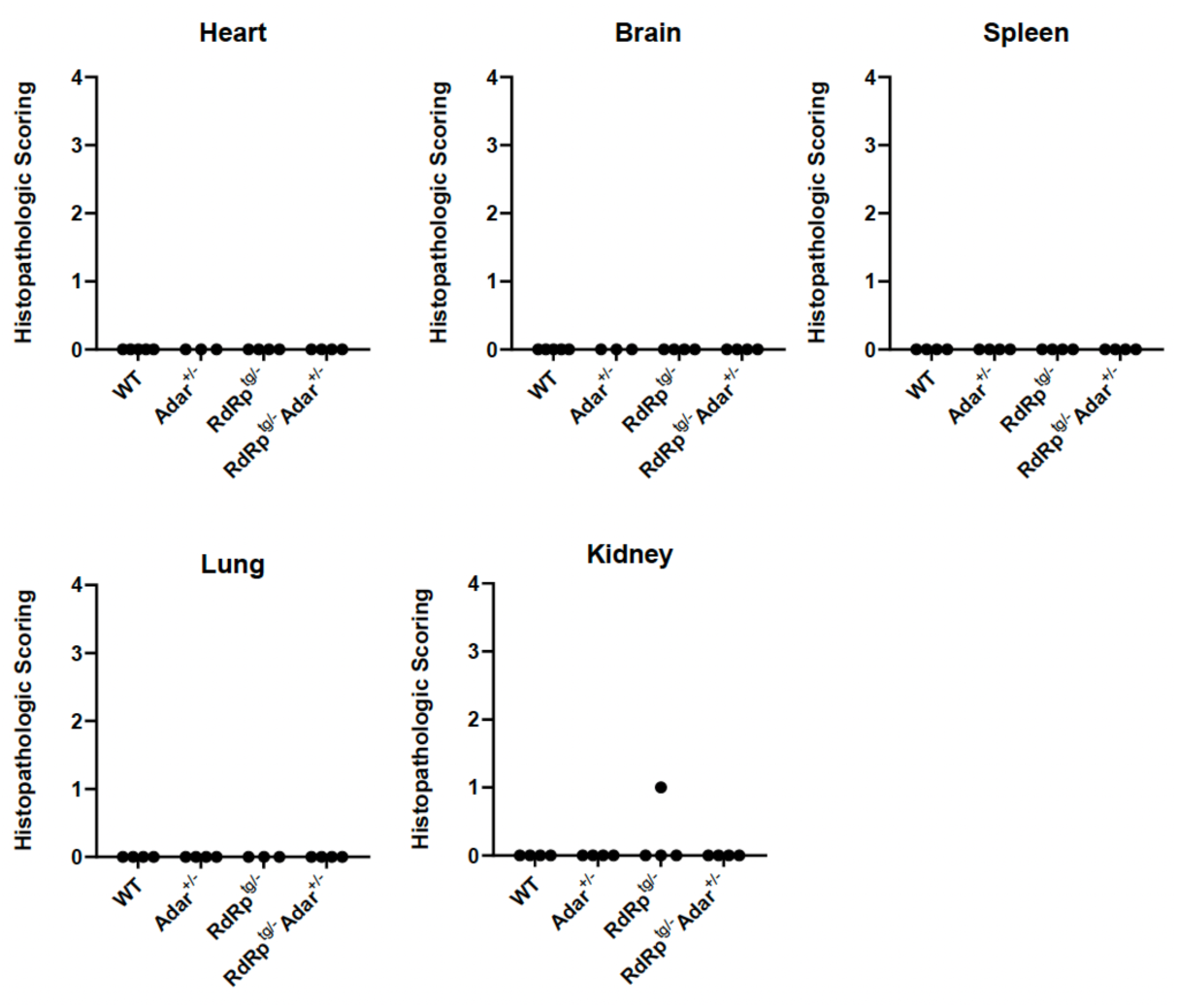
Histopathological grading of mouse tissues. No significant changes or differences were noted in the RdRp^tg/-^ Adar^+/-^ animals to explain the clinical presentations of this genotype.

